# Non-centromeric CENP-A epigenetically regulates epithelial-mesenchymal plasticity and heterogeneity in human cells

**DOI:** 10.1101/2025.09.09.672651

**Authors:** Charlène Renaud-Pageot, Daniele Capocefalo, Sébastien Lemaire, Audrey Forest, Laura Cantini, Geneviève Almouzni

**Author notes:** These authors contributed equally (Daniele Capocefalo, Sébastien Lemaire).

## Abstract

The centromeric histone variant CENP-A, highly expressed in aggressive cancers, can promote an epithelial-mesenchymal transition (EMT) whose underlying mechanisms remain to decipher. Here, we tracked both the temporal dynamics of EMT states and CENP-A localization over time using a reversible high CENP-A expression system in human cells. Cell populations presenting hybrid EMT states at start, when exposed to high CENP-A levels, progressively accumulated mesenchymal states and displayed increased centromeric and ectopic CENP-A incorporation. Mechanistically, we reveal ectopic CENP-A gains at EMT genes by chromatin immunoprecipitation and identify two distinct EMT programs activated at different stages of the cell cycle by single-nucleus multi-omics. Importantly, while a pre-existent inflammatory program got amplified, high CENP-A induced a new developmental program. Remarkably, interrupting high CENP-A provision erased induced programs along with ectopic CENP-A incorporation, in line with non-genetic alterations. Our findings uncover an unconventional, non-centromeric function for CENP-A in epigenetically modulating epithelial-mesenchymal plasticity.

## Introduction

The Centromere Protein A (CENP-A) is a histone H3 variant specifically incorporated into centromeric chromatin and essential for the proper segregation of human chromosomes to daughter cells^1–3^. CENP-A combines both genetic and epigenetic features to provide the foundation of kinetochore assembly during cell division^4^, and to determine^5,6^ and maintain^7–10^ centromere position across mitotic and meiotic divisions. CENP-A restriction to the centromere depends on key parameters. First, the exclusive partnership between CENP-A and its dedicated histone chaperone, the Holiday Junction Recognition Protein (HJURP), ensures the recycling of parental CENP-A at the replication fork^11^ and directs the specific incorporation of newly synthetized CENP-A at centromeres in late mitosis^12,13^. Second, the partnership between CENP-A and HJURP is reinforced by co-regulating their levels during the cell cycle with a peak of expression in G2/M^14^. They both harbor in their promoters a CDE/CHR motif which is recognized by the DREAM complex^15^, enabling to mediate a p53-dependent transcriptional repression^18^. Third, chromatin replication acts as an error correction mechanism removing CENP-A from chromosome arms, keeping CENP-A nucleosomes specifically at centromeres^17^.

Elevated CENP-A expression is frequently observed across diverse cancer types and can reach over 1000-fold higher levels relative to healthy tissues^18^. This upregulation occurs in both p53-defective and wild type contexts, as CENP-A can evade p53-mediated regulation^19,20^. Increased CENP-A levels disrupt the balance in the proportions of histone H3 variants, and the excess of CENP-A hijacks H3.3-dedicated chaperones DAXX^21^ and HIRA^22^. This imbalance enables the misincorporation of CENP-A in non-centromeric chromatin, including intergenic and promoter regions^21–27^. Ectopic CENP-A traps a series of kinetochore components away from native centromeres^28–30^, likely weakening centromere function, and promoting chromosomal instability and aneuploidy^20,29,30^. While high CENP-A levels may reflect the increased mitotic activity of cancer cells, it may also represent an active process participating to cancer progression and therapeutic resistance. We previously reported that high CENP-A levels could promote an epithelial-mesenchymal transition (EMT) in p53-defective cells^19^. EMT is a major cell fate transition during which cells switch from an epithelial to a mesenchymal state, a phenotypic plasticity associated with cancer progression, resistance and metastasis^31^. In line with these observations, high CENP-A levels correlate with cancer aggressiveness, invasion and metastasis across multiple tumor types^18^. Importantly, EMT is not a single program but occurs through distinct pathways driving either an inflammatory EMT which contributes to adult wound healing and fibrosis^32,33^, or a developmental EMT which governs embryonic cell migration and morphogenesis^34,35^. However, tumor cells can hijack both programs to implement inflammation and developmental cell migration^36^. Furthermore, since formulation of the concept^37,38^, increasing evidence has supported the existence of hybrid (or intermediate) states between epithelial and mesenchymal in the context of fibrosis^39,40^, development^41^, and cancer in circulating tumor cells^42–44^ and mouse models^45^. Thus, the spectrum of states observed in a given cellular context has been defined as epithelial-mesenchymal plasticity (EMP), a means to reflect the ability of cells to interconvert between different states^31^. Accordingly, high phenotypic plasticity and diversity could allow progression toward a mesenchymal fate through multiple alternative paths. However, the dynamics of these hybrid states and the molecular actors governing cancer-associated EMT states remain unresolved. Thus, understanding how CENP-A promotes EMT, particularly whether it impacts plasticity, with the potential to use multiple paths and/or to select specific EMT programs, is relevant in the context of cancer and normal development.

In this study, we wished to determine how this key centromeric protein could promote EMT, which pathways were involved, whether it required a critical time window, and whether it was reversible. To this aim, we monitored the temporal dynamics of EMT states in a model of fibrocystic human breast cell line conditionally expressing elevated CENP-A levels. We characterized specific transition states by combining microscopy and flow cytometry. We found that the induction of high CENP-A levels promotes EMT at the expense of cells displaying a hybrid state. We next identified the transition programs specifically elicited by transcriptomics. Mechanistically, we revealed ectopic CENP-A gains at EMT gene loci by chromatin immunoprecipitation followed by sequencing (ChIP-seq), accompanied by the activation of two distinct EMT programs at different stages of the cell cycle, as determined by single cell multiomics combining ATAC and RNA sequencing. Finally, by interrupting CENP-A induction, we could erase the ectopic CENP-A from chromosome arms along with induced mesenchymal states. Thus, while CENP-A has long been considered for its key role in centromere function and genome stability, we unveil here a non-genetically linked feature of CENP-A in regulating epithelial-mesenchymal plasticity. We discuss this unanticipated epigenetic role for CENP-A outside centromere and its implications in cancer.

## Results

### High CENP-A levels promote mesenchymal states in cell lines with hybrid EMT states

To identify the features enabling high CENP-A levels to promote a mesenchymal phenotype, we used a previously characterized doxycycline-inducible system^19^ (Tet-On–CENPA–FLAG–HA; see Material and Methods) in several human cell lines (Table 1; Figure S1A). We performed co-immunostaining for epithelial markers (E-cadherin or β-Catenin, forming a complex at the membrane of epithelial cells^46^) and a mesenchymal marker (Vimentin) to identify EMT states (Figure 1A). In control conditions, we confirmed the detection of epithelial and mesenchymal populations in both p53 wild type (WT; transduced with an empty vector) and p53-defective (transduced with a constitutively expressed dominant-negative TP53 construct) MCF10-2A cells, derived from breast tissue with benign fibrosis^47^ (Figure 1B). Strikingly, we identified for the first time a co-existing population of cells in a hybrid EMT state, characterized by the co-expression of epithelial and mesenchymal markers, present in both p53 backgrounds (Figure 1B). To characterize the temporal dynamics of EMT progression, we conducted time course experiments over a 35-day period, monitoring the distribution of cell states before, during and after EMT (Figures 1A and S1A-C). Without CENP-A induction, the epithelial population increased (from 50 % to 80%) in both p53-WT and p53-defective cells, while the mesenchymal population remained low (< 4%), and the hybrid population declined from 45% to ∼10% (Figures 1C and S1E). However, induction of high CENP-A expression altered these proportions in the p53-defective cells. Indeed, the epithelial population remained stable, hybrid decreased (from 45% to ∼10%), and mesenchymal population increased from day 10 onwards to stabilize by day 20 (at ∼30%) (Figure 1C), a sigmoidal pattern consistent with theoretical models of EMT progression^48–51^. Importantly, our flow cytometry analysis confirmed these results (Figure S1D). Notably, upon CENP-A induction, hybrid cells showed variability in the EMT spectrum, with more advanced states exhibiting high vimentin levels throughout their cytoplasm, generally at the periphery of cell colonies (Figure S1C), as illustrated by quantifying vimentin intensity relative to control (Figure 1D). In p53-WT cells, the mesenchymal accumulation was not comparable (Figure S1E), likely due to CENP-A-induced cell cycle arrest^19^. Our cell culture medium containing epidermal growth factor (EGF), an EMT inducer^52–54^, we tested whether a simple variation in EGF concentrations could promote or potentiate EMT in p53-defective cells. In the absence of EGF, with reduced cell proliferation^55^, only 1% of cells harbored a hybrid state and these conditions abolished the capacity of CENP-A induction to promote the mesenchymal phenotype (Figure S1F). Conversely, increasing EGF concentration up to 20X was not sufficient to promote hybrid nor mesenchymal (remaining at ∼4%) states (Figure S1F), indicating that enhanced EGF signaling cannot recapitulate the effects of CENP-A induction. Finally, we examined whether the impact of CENP-A induction depends on the existing populations of EMT states at start across different cell lines. In cell lines where we did not detect mesenchymal cells upon CENP-A induction (HCC1954^19^, MCF7 and T47D), we only found epithelial cells present at start (Figure S1G, top). Remarkably, we found that the commonly used, near-diploid RPE-1 cell line displayed an advanced hybrid EMT phenotype, and CENP-A induction triggered a switch to a complete mesenchymal state within 10 days (Figure S1G, bottom). These findings support two main conclusions concerning how CENP-A can induce EMT. First, that it occurs in cell lines in which we identified hybrid EMT states, and second, that the presence of these hybrid states diminishes while mesenchymal states increase, prompting us to consider altered cell population dynamics.

**Figure 1.**
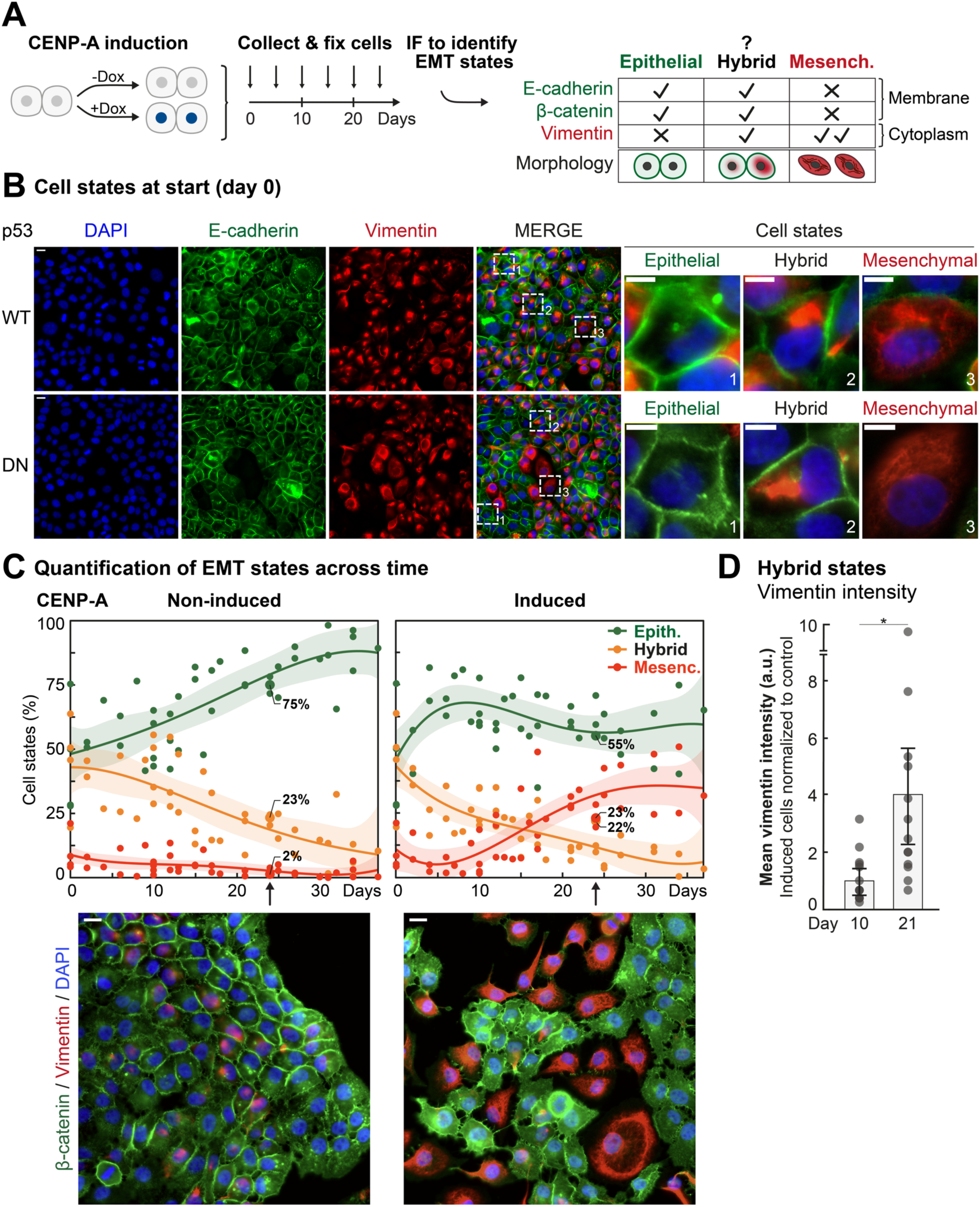
Impact of chronically elevated CENP-A levels on EMT states. (A) Experimental strategy. From the same starting population (day 0), we split cells into non-induced (-Dox, control) or induced for high CENP-A expression (+Dox) and collected them for fixation at various time points until day 37. We used immunofluorescence (IF) microscopy to detect epithelial, hybrid, and mesenchymal states using specific antibodies as indicated. (B) EMT states at day 0 in the MCF10-2A cell line. *Left:* Representative epifluorescence images of cells immunostained for E-cadherin (green, epithelial) and vimentin (red, mesenchymal), with DNA counterstained with DAPI (blue). Scale bars = 20 µm. Squared boxes indicate cells on the right. *Right:* Insets of representative cell states. (1) Epithelial: green signal at the membrane, little or no red signal, polygonal cell; (2) Hybrid: green signal at the membrane with an intense red speckle which can spread out in the cytoplasm, polygonal or round cell; (3) Mesenchymal: loss of epithelial marker at the membrane, red marker extending throughout the cytoplasm as a network of filaments, round or spindle-shaped cell. Scale bars = 10 µm. (C) EMT states across time in the MCF10-2A cell line. *Top:* Proportions of epithelial (green), hybrid (orange) and mesenchymal (red) states. Plots show the trendline as polynomial of the fourth degree ± 95% confidence interval (shaded area). n > 200 cells across ≥ 2 fields for each data point. N ≥ 5 biologically independent experiments. Numbers indicate proportions of states in the images below. *Bottom:* Representative epifluorescence images of cells 24 days post-induction. β-catenin (green), vimentin (red), and DAPI (blue). Corresponding CENP-A stainings in Figure S1C. (D) Mean vimentin intensity in hybrid cells at days 10 and 21 post-induction (relative to non-induced control). Plots show mean signal in hybrid cells ± 95% confidence interval. Dots represent mean per microscopic field. N = 3 biologically independent experiments. Based on IF analysis, image segmentation depicted in Figure S1. Statistical significance tested by two-tailed Welch’s t-test. * = p-value < 0.05.

**Table 1.** Description of cell lines used in the study.

| Cell line | Sex | p53 status | Cell type | Comments |
| --- | --- | --- | --- | --- |
| RPE-1 | Female | Wild-type<sup>a</sup> | Eye (retina), non-tumoral | hTERT immortalized |
| MCF10-2A | Female | Wild-type<sup>a</sup> | Breast, non-tumoral, fibrocystic | Spontaneously immortalized |
| MCF7 | Female | Wild-type | Breast, invasive carcinoma | HER- ER+ PR+ luminal |
| T47-D | Female | LOF mutation (L194F) | Breast, invasive carcinoma | HER- ER+ PR+ luminal |
| HCC19-54 | Female | LOF mutation (Y163C) | Breast, invasive carcinoma | HER+ ER- PR- luminal |

<sup>a</sup>p53-defective cells also used (transduction with dominant negative TP53 construct)

### Distinct mechanisms promote mesenchymal identity beyond DNA damage and proliferation advantage

To further investigate how high CENP-A levels could promote mesenchymal states at the expense of hybrid states, we started with assessing its impact on centromere function. We evaluated two potential mechanisms (Figure 2A): (**1**) Increased CENP-A levels promote mitotic defects, leading to chromosomal instability (CIN)^29,30^ and potential DNA damage^56^. Both mechanisms could in turn favor EMT, as observed in cancer cells^57–59^. (**2**) Altered rates of cell death or division could shift the relative abundance of cell states in the population. First, to examine CIN, we quantified micronuclei^60^ and other nuclear alterations, including envelope rupture^60^, deformation^61^, multinucleation^62^ and mitoses with defective chromosome segregation, across EMT states when mesenchymal states increase (days 10-20) (Figure 2B, left). Our analysis confirmed that high CENP-A levels lead to increased mitotic defects in all states and, relative to other states, the mesenchymal population did not display a higher increase in CIN (Figure 2B, right). To look into possible DNA damage response, while we and others previously reported that high CENP-A levels did not lead to significant increase in acute DNA damage or in the rate of DNA repair^19,20^, here, we examined longer timescales matching our time course experiments. We found that γ-H2AX levels, an early marker of DNA damage, did not significantly increase and could instead be higher after returning to basal CENP-A levels (Figure S2A). Second, we examined how CENP-A induction may affect cell survival, considering that loss of HJURP and CENP-A promotes apoptosis in p53-null transformed mouse cells^16^. During the same time window as our prior experiments, we used immunofluorescence (IF) staining for cleaved caspase 3, an early marker of apoptosis (Figure 2C, left). In the control condition, mesenchymal cells exhibited a significantly higher signal compared to other populations (∼40% mesenchymal, ∼10% hybrid and <2% epithelial cells positive for the marker). However, this was partially reduced upon CENP-A induction, with a ∼10 to 15% decrease in both hybrid and mesenchymal cells (Figure 2C, right). Although reduced apoptosis may facilitate mesenchymal expansion, this limited effect was insufficient to explain the observed phenotype (Table 2), raising the possibility of additional mechanisms involved. We then assessed whether proliferation advantage could contribute to the relative increase in mesenchymal cells. We tracked cell divisions using carboxyfluorescein diacetate succinimidyl ester (CFSE), a fluorescent dye that is equally distributed between daughter cells at each division. We thus measured signal dilution as a proxy to evaluate the extent to which a cell (identified as epithelial, hybrid or mesenchymal) has divided. The assay was carried out between day 12 and day 16, when mesenchymal cells increase in proportion (Figure 2D, left). Strikingly, CENP-A induction led to a reduced proliferation rate across all populations (Figure 2D, right). Mesenchymal cells exhibited the slowest proliferation rate while hybrid states showed the fastest, arguing against a proliferation advantage as the main driver of the mesenchymal phenotype. Simulation based on the estimated doubling time and cell death rate failed to account for the ∼25% increase in mesenchymal cells (Table 2). Taken together, the observed increase could not suffice to provide a convincing argument for a role attributed solely to DNA damage, differences in cell death, or proliferation, leading us to examine whether transcription switches could operate over time.

**Figure 2.**
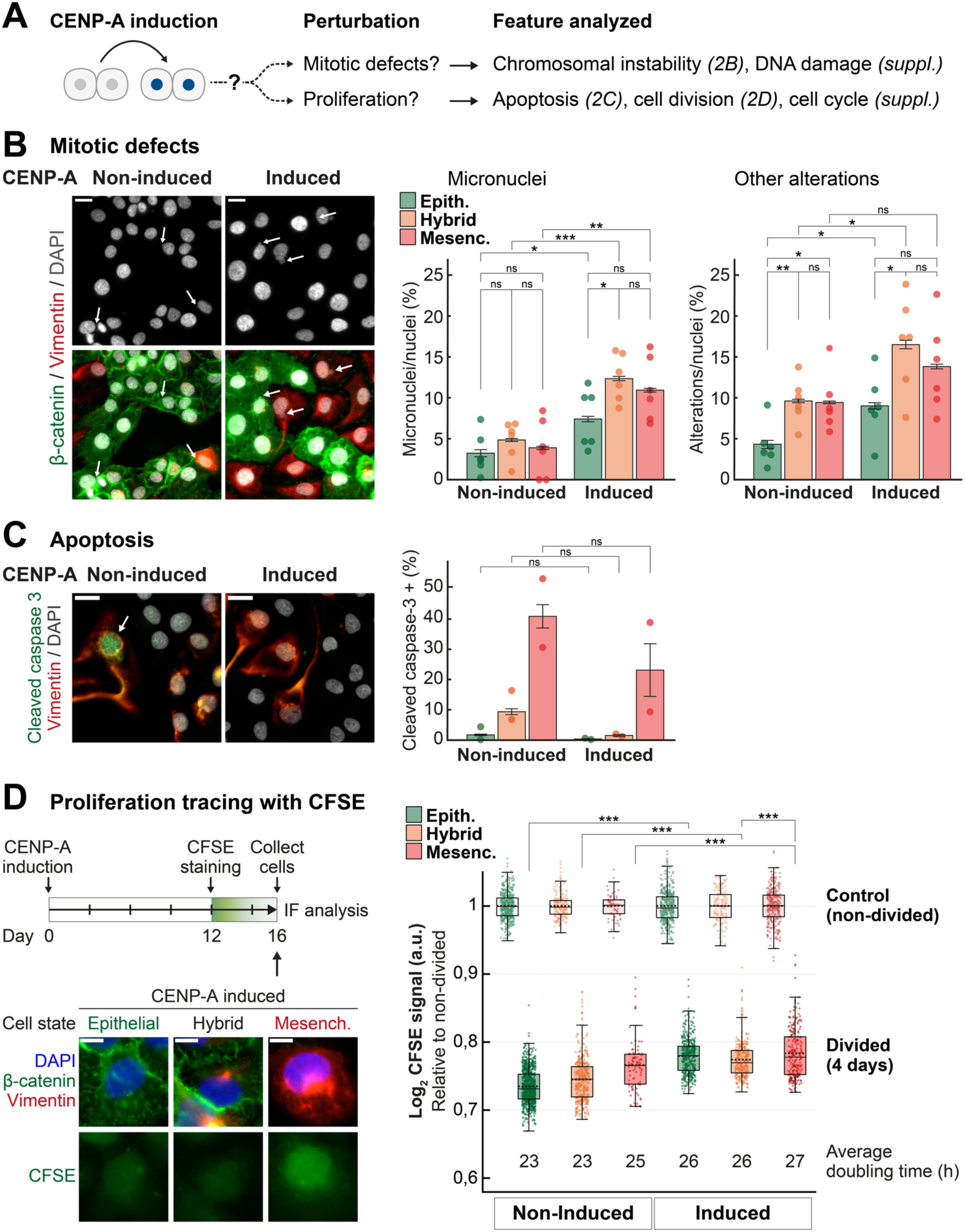
Analysis of mitotic defects and proliferation in promoting EMT. (A) Scheme for the experimental rationale to assess the impact of high CENP-A levels on mitotic defects and proliferation in MCF10-2A cells, and their contribution in promoting mesenchymal EMT states. (B) Mitotic defects. *Left:* Epifluorescence images at day 15 showing β-catenin (green), vimentin (red), and DAPI (grey). Arrows denote micronuclei and segregation defects. Scale bars = 20 µm. *Right:* Quantification of micronuclei and other nuclear alterations (deformation, envelope rupture, multinucleation and chromosome mis-segregation events) for each EMT state between days 12 and 17. Plots show mean ± 95% confidence interval. Dots indicate mean per experiment. n > 400 nuclei/state across ≥ 3 fields for each replicate. N = 6 biologically independent experiments. Statistical significance tested by two-tailed Welch’s t-test. * = p-value < 0.05, ** = p-value < 0.01, *** = p-value < 0.001. (C) Apoptosis. *Left:* Epifluorescence images at day 15 showing cleaved (active) effector caspase-3 (green), vimentin (red), and DAPI (grey). Scale bars = 20 µm. *Right:* Proportions of cells with positive cleaved caspase-3 signal between days 15 and 24. Plots show mean ± 95% confidence interval. Dots indicate mean per experiment. n > 150 nuclei/state across ≥ 3 fields for each replicate. N = 2 independent experiments. Statistical significance tested by two-tailed Welch’s t-test. (D) Cell proliferation. *Top left:* Experimental scheme. We cultivated cells-Dox or +Dox. At day 14, we pulse-labeled one plate per condition with CFSE (test). At day 16, we pulse-labeled a second plate per condition using the same CFSE solution (undivided control). We harvested all plates and immunostained cells for EMT markers. We measured CFSE nuclear intensity per EMT state. *Bottom left:* Representative epifluorescence images of EMT states showing β-catenin (green), vimentin (red), DAPI (blue) and CFSE (green, lower panel). *Right:* Log_2_-transformed CFSE intensity in divided *versus* control cells. Plots show mean (solid line), median (dotted line) and standard deviation for 2 biologically independent experiments. Each dot represents one cell. n > 600 cells per condition. Numbers indicate the estimated doubling times based on average signal dilution (see Methods). Statistical significance tested by two-tailed Welch’s t-test. * = p-value < 0.05, ** = p-value < 0.01, *** = p-value < 0.001.

**Table 2.** Percentages of EMT states observed *versus* simulated at day 16.

|  | Non-induced |  |  |  | Induced |  |  |
| --- | --- | --- | --- | --- | --- | --- | --- |
|  | d12 | d16<br>observed | d16<sup>a</sup><br>simulated |  | d12 | d16<br>observed | d16<sup>a</sup><br>simulated |
| % Epithelial | 58,5 | **48** | **66** |  | 58,5 | **55** | **62** |
| % Hybrid | 30,5 | **41** | **28** |  | 27,5 | **24** | **28** |
| % Mesenchymal | 11 | **11** | **5** |  | 14 | **21** | **10** |

<sup>a</sup> $(N_i \cdot 2^{(96/T)} - D)/N_T$
Considering the number of cells seeded at d12 ( $N_i$ ), the total cell count at d16 ( $N_T$ ), the estimated doubling time ( $T$ ) and the fraction of apoptotic cells ( $D$ ).

### Elevated CENP-A expression promotes two EMT programs associated with CENP-A enrichment

To explore how CENP-A induction induces the mesenchymal phenotype in our cell population, we examined the impact on transcription with a particular focus on EMT signaling pathways. To this aim, we prepared RNA focusing on days 10 and 24, two time points prior and after detection of significant changes (Figure 1C). Bulk RNA-seq data analysis enabled to identify distinct EMT effectors, corresponding respectively to the inflammatory and the developmental EMT programs (Figures 3A and S3A). First, we examined the situation with respect to p53 status and found that p53 inactivation alone did not significantly impact EMT-related transcriptional programs at the bulk level (Figure 3A, top). Second, in a p53-WT background, CENP-A induction on its own favored a distinct transcriptional signature characterized by upregulation of genes within the inflammatory EMT axis (e.g., *IL6, IL1B, TGFB2*). This was accompanied by a downregulation of genes associated with the developmental axis (e.g., *AKT3*, *CTNNB1*, *FZD2/4*) (Figure 3A, middle). In a p53-defective context, not only CENP-A induction further enhanced the inflammatory EMT response (*IL6*, *JAK, STAT*) but also induced developmental signaling pathways (*AKT3*, *CTNNB1*, *FZD2/4, TCF7L1, WNT3*). Interestingly, this latter induction occurred without engaging *TGFB2*, which was instead significantly repressed (Figure 3A, bottom). Thus, CENP-A induction engages both inflammatory and developmental EMT axes, with the latter specifically induced in p53-defective cells. Notably, in a second, biologically independent experiment, we detected the activation of the same pathways as early as day 10 (prior to the increase in mesenchymal states), with further enhancement by day 24 (Figure S3A). We also confirmed at day 1 post-induction the requirement for chronic induction to elicit these changes (Figure S3B). We then wished to validate these effects at the protein level and assessed IL-6 expression and β-catenin activation through its nuclear translocation. Western blot analyses revealed increased IL-6 levels (Figure 3B) and elevated levels of non-phosphorylated (stable/active) β-catenin (Figure S3C), while IF analysis showed β-catenin translocation (Figure 3C), confirming the impact of CENP-A on these pathways. Next, to investigate whether elevated CENP-A expression levels lead to local CENP-A enrichment at specific gene loci, we carried out ChIP-seq experiments at the same time points (day 10 and 24). This way, we could compare our transcriptomic data with the ChIP-seq profiles corresponding to CENP-A and H3.3 for a genome-wide analysis (Figures 3D, S3D, S3E). Based on our previous findings^21^, we expected to detect CENP-A at sites previously marked with H3.3, including high turnover regions and transcription start sites. Here, the combination of ChIP enabled to directly confirm CENP-A and H3.3 co-enrichment (Figure S3D). Furthermore, among the genes linked to both development and inflammation programs, we detected an early gain in CENP-A prior to the increase in mesenchymal cells (by day 10). This enrichment persisted at day 24 (Figure 3D, top left). Our gene ontology (GO) enrichment analysis (Figure 3D, bottom) confirmed that the corresponding genes fall into the same pathways detected in our bulk RNA-seq (*JAK/STAT*, *AKT*, *Wnt*). Among genes described above, the early gain in CENP-A (Figure 3D, top right) corresponded to genes affected by CENP-A induction (Figure 3A). Based on these results, we propose that high CENP-A expression allows its stable incorporation in chromatin at genes encoding EMT factors. This in turn could favor a chromatin plasticity enabling the activation of distinct EMT gene expression programs.

**Figure 3.**
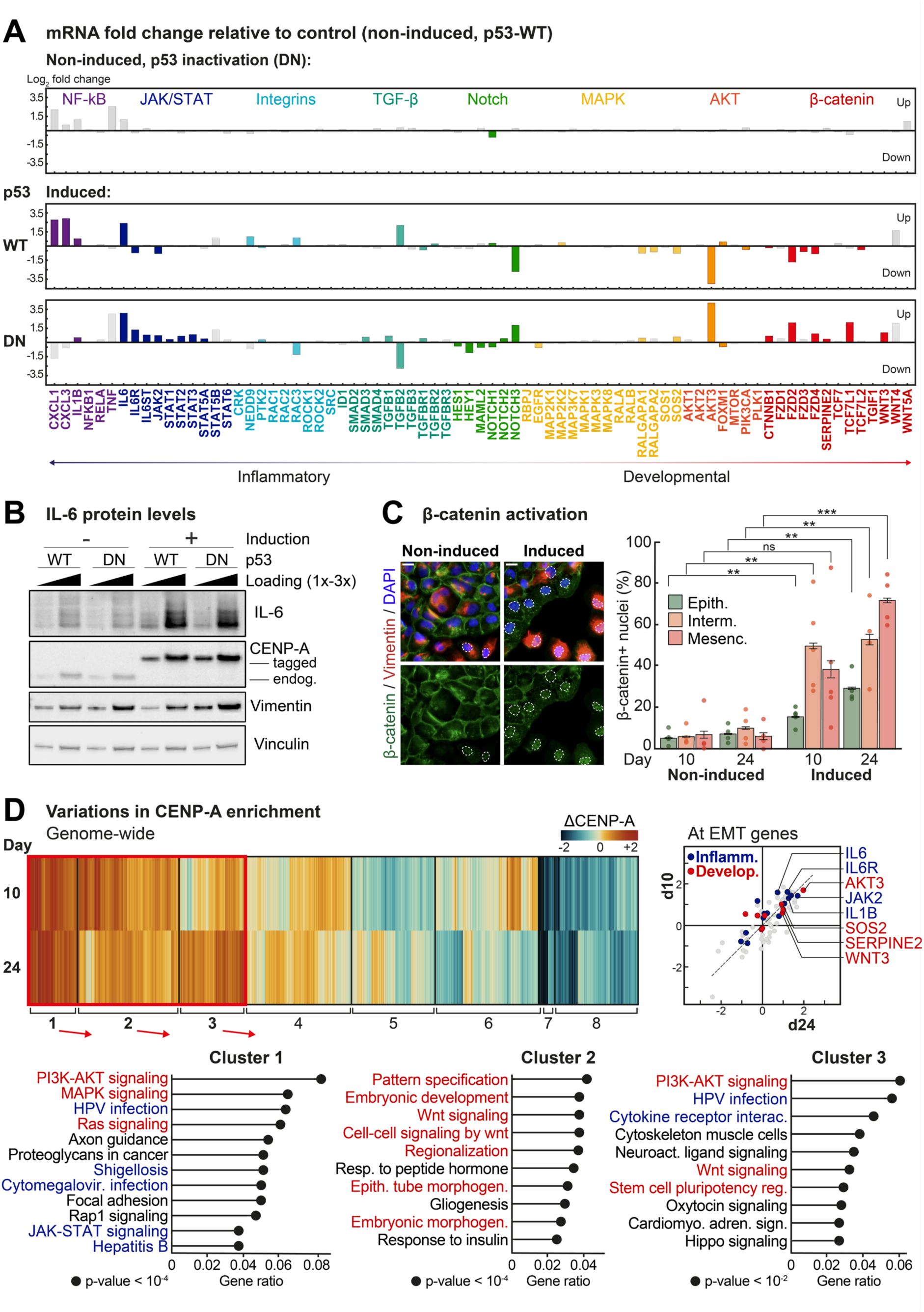
EMT programs impacted by increased CENP-A levels and CENP-A mislocalization. (A) mRNA expression analysis of EMT genes in MCF10-2A cells at day 24. Bar plots show log_2_ fold change relative to the p53-WT non-induced condition (TMM-normalized). Genes are grouped by signaling pathway (indicated above) and functional category (inflammation or development). Colored bars indicate statistically significant changes (adjusted p-value < 0.05). (B) Western blot analysis of total protein extracts from MCF10-2A cells (-Dox or +Dox), with vinculin as loading control. Primary antibodies are indicated on the right. *IL-6 detection as multiple bands, consistent with its presence as differentially modified (glycosylated) isoforms, with apparent molecular weights from 23 to 30 kDa. (C) β-catenin localization in p53-DN MCF10-2A cells at days 10 and 24. *Left:* Epifluorescence images showing β-catenin (green), vimentin (red) and DAPI (blue). Scale bars = 40 µm. *Right:* Proportions of cells with positive β-catenin nuclear signal. Plots show mean ± 95% confidence interval for 4 or 5 independent experiments. Dots indicate mean per experiment. n ≥ 600 cells/state across ≥ 3 fields for each replicate. Statistical significance tested by two-tailed Welch’s t test. * = p-value < 0.05, ** = p-value < 0.01, *** = p-value < 0.001. (D) ChIP-seq analysis of CENP-A at transcription start sites (TSS) on days 10 and 24 post-induction (relative to non-induced). *Top left:* Heatmap of CENP-A enrichment at gene TSS. Columns represent protein-coding genes, clustered by changes in CENP-A and H3.3 occupancy at their TSS. Rows indicate relative gain or loss of CENP-A (log_2_-transformed, input-normalized, sequencing depth-normalized, and mean-centered). Corresponding H3.3 heatmap shown in Figure S3. *Bottom left:* Gene ontology (GO) enrichment analysis for three gene clusters. Resp., Response; Cardiomyo. Adren. Sign., Cardiomyocyte adrenergic signaling; Neuroact., Neuroactive. *Right:* Relative gain in CENP-A at EMT gene TSS. Colored dots indicate genes significantly upregulated upon CENP-A induction (from panel A).

### The two CENP-A-induced EMT programs follow distinct trajectories linked to cell cycle as discriminated by single-nucleus multi-omics

We next explored at a single cell resolution how CENP-A promotes the inflammatory and developmental EMT axes considering both transcription programs and cell cycle. Since our microscopy and cytometry analyses revealed a heterogeneous and dynamic population, we carried out single cell resolution analysis at days 10 and 24 (prior and after mesenchymal increase). At these time points, for the same nuclei we jointly profiled single-nucleus RNA (snRNA) and ATAC (snATAC) data using 10X Multiome (Figure 4A). First, we observed a global increase in chromatin accessibility upon CENP-A induction (Figure S4A), in line with an increased plasticity enabling the activation of distinct EMT gene expression programs as we proposed. We then integrated the readings from both modalities (snRNA and snATAC) using Mowgli, a matrix factorization method defining lower-dimensional cell embeddings from paired single-nucleus data^63–65^ (see Methods) (Figures 4A and S4B). While we could readily distinguish mesenchymal and non-mesenchymal populations (Figure 4B), we could not isolate the hybrid population (cells expressing both epithelial and mesenchymal markers, Figure 4B) from the epithelial cells. However, we could monitor changes in cell identity since CENP-A induction significantly increased the proportion of mesenchymal cells over time (Figure 4B). Thus, we could identify the specific key cellular pathways driving EMT in our conditions. To identify co-regulated subsets of differentially expressed genes (DEGs), we applied Leiden clustering to the multi-omics cell embedding from Mowgli and revealed two mesenchymal clusters termed “7” and “12” (Figures 4C and S4C). Within these clusters, we classified cells according to their position in the cell cycle (G2/M, S and G1 phases) (Figure 4D). For cluster 7, spontaneous and induced mesenchymal cells occurred in G1 phase or cell cycle exit whereas, for cluster 12, induced mesenchymal cells occurred predominantly in S and G2/M phases. Thus, high CENP-A could induce EMT in all cell cycle phases (clusters 7 and 12) (Figure 4C-D), as confirmed by FACS (Figure S4D). To further study cell dynamics, we inferred RNA velocity using scVelo^66^ on the multi-omics embedding (see Methods). In control cells, a single EMT trajectory led to mesenchymal cluster 7 (Figure 4E). However, CENP-A induction resulted in two trajectories: a first one, as in control cells, directed toward cluster 7, and a new one, unique to this condition, toward cluster 12 (Figure 4E). In both clusters, top DEGs included key EMT drivers such as RBMS3 (Figure S5A,B), an RNA-binding protein that stabilizes mRNAs encoding the EMT transcription factor PRRX1^67^. Cluster 7 associated with high expression of inflammation factors including fibrosis drivers (IL6, HIPK2^68^, NEAT1^69^) (Figure S5B, middle) and enrichment for genes regulating cell division and apoptosis (GO analysis relative to other clusters; Figure 4C, bottom). In contrast, cluster 12 displayed high expression of proliferation markers (Figure S5A, B) and enrichment for developmental and Wnt signaling pathways (GO analysis; Figure 4C, bottom). Notably, developmental factors specifically emerged in cluster 12, while inflammation factors distributed broadly across other clusters, reflecting the fibrocystic origin of MCF10-2A cells^47^. Furthermore, to infer candidate transcriptional regulators involved in these distinct EMTs, we applied HuMMuS^70^, a method using both snRNA and snATAC modalities to reconstruct gene regulatory networks. This analysis revealed, for cluster 7, two groups of strongly reduced activities involved in S-phase progression (CREB1-JUN and ETV6-ETV4-ETV1-ETS1) and, for cluster 12, increased activity of regulons upstream of Wnt and AKT signaling, linked to embryonic development (e.g., pioneer factor TFAP2, required for neural crest induction^71^) (Figure S5C). Thus, control cells used the inflammatory EMT that only occurs in G1 phase or outside the cell cycle. By contrast, upon CENP-A induction, two specific trajectories emerged at different cell cycle stages. Thus, the CENP-A-induced programs detected in our bulk RNA and ChIP-seq data (Figures 3A-D) followed distinct, cell cycle-dependent trajectories. We conclude that CENP-A, by increasing chromatin plasticity, broadens the spectrum of available EMT trajectories in hybrid cells at different times during the cell cycle.

**Figure 4.**
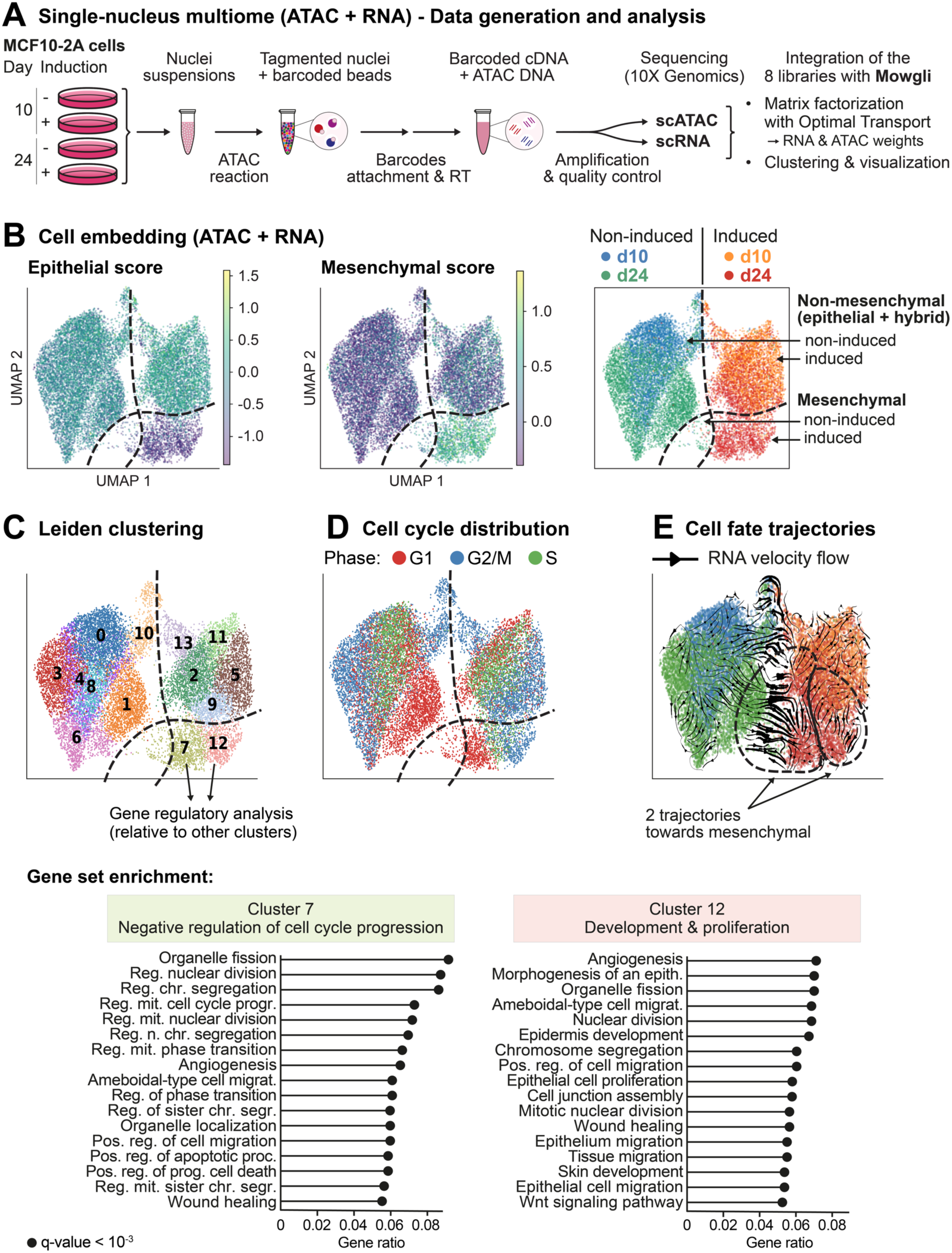
Single-nucleus multi-omic profiling of EMT programs and trajectories. (A) Experimental workflow to profile snRNA-seq and snATAC-seq simultaneously within the same cells. We collected p53-DN MCF10-2A cells at days 10 and 24 post-induction (-Dox and +Dox). Isolated nuclei were tagmented, then associated with gel beads containing a poly(dT) sequence (for production of barcoded cDNAs) and a Spacer sequence (for barcode attachment to transposed DNA fragments). Incubation of the GEMs and reverse transcription (RT) produced 10x barcoded DNA from the transposed DNA and 10x barcoded cDNA from poly-adenylated mRNAs, that were sequenced with 10X Chromium Multiome. Data integration was performed using Mowgli (Non-negative Matrix Factorization with Optimal Transport). (B) Uniform Manifold Approximation and Projection maps (UMAPs) of Mowgli embeddings. *Left and middle:* Epithelial and mesenchymal scores based on CDH1 and EPCAM (epithelial), and VIM, FN1 and ZEB1 (mesenchymal) expression. *Right:* Mesenchymal and non-mesenchymal subpopulations among the samples: non-induced at days 10 (blue) and 24 (green) and induced at days 10 (orange) and 24 (red). (C) *Top:* UMAP with Leiden clustering applied to Mowgli embeddings. *Bottom:* GO enrichment analysis of DEGs in clusters 7 and 12, relative to the other clusters. Reg. nuclear division, Regulation of nuclear division; Reg. chr. segregation; Regulation of chromosome segregation; Reg. mit. cell cycle progr., Regulation of mitotic cell cycle progression; Reg. mit. nuclear division, Regulation of mitotic nuclear division; Reg. n. chr. segregation, Regulation of nuclear chromosome segregation; Reg. mit. phase transition, Regulation of mitotic cell cycle phase transition; Ameboidal-type cell migrat., Ameboidal-type cell migration; Reg. of phase transition, Regulation of cell cycle phase transition; Reg. of sister chr. segr., Regulation of sister chromatid segregation; Pos. reg. of cell migration, Positive regulation of cell migration; Pos. reg. of apoptotic proc., Positive regulation of apoptotic processes; Pos. ref. of prog. cell death, Positive regulation of programmed cell death; Reg. mit. sister chr. segr., Regulation of mitotic sister chromatid segregation; Morphogenesis of an epith., Morphogenesis of an epithelium; Pos. reg. of cell migration, Positive regulation of cell migration. (D) UMAP showing inferred cell cycle phases. (E) Cell fate trajectories inferred by RNA velocity. The streamline plot projected onto the UMAP depicts the dynamical modelling. Arrows (gene-averaged velocity vectors) indicate the main directional flow of cell states over time.

### CENP-A is an epigenetic driver of EMT requiring its ectopic localization

To further evaluate whether our mesenchymal population had acquired stable genetic alterations, we tested whether interrupting the induction to revert CENP-A to basal levels could rebalance cell populations. We induced high CENP-A expression for 24 days, then, by removing doxycycline from our cell culture media, we returned to basal levels and followed cell states over 10 days (Figure 5A, top). We monitored the proportions of cells in each state (epithelial, hybrid or mesenchymal) in comparison to non-reversed cells, alongside CENP-A subnuclear localization (Figure 5A, top right) as we found distinct CENP-A patterns in interphase nuclei (Figure S1A). Indeed, in control cells, as previously documented^17,19^, our staining showed CENP-A in small foci at centromeres co-localizing with CENP-B (Figure S1A, top). Upon CENP-A induction, these foci enlarged and CENP-A signal increased as a strong diffuse pattern, indicative of ectopic (non-centromeric) localization (Figure S1A, bottom). Interestingly, by interrupting CENP-A induction, we removed the diffuse CENP-A signal while preserving large CENP-A foci corresponding to centromeres (Figures 5A, 5C and S6A). Remarkably, this CENP-A marking in large foci enabled to recognize cell populations that had previously experienced high CENP-A levels. We found that, after 10 days of interruption, the mesenchymal population decreased from 30% to less than 10% on average, as measured by both microscopy and flow cytometry (Figures 5B and S6B). Importantly, the mesenchymal population decrease did not correspond to an increase in the hybrid population, but rather in the epithelial population (Figures 5B and S6B) to reach the proportions observed in control cells. Yet, cells showed a clear change in mesenchymal cell morphology (Figures 5A and S6C), with decreased levels of translocated and non-phosphorylated (stable/active form) β-catenin (Figures 5D, S6C and S6D) as well as decreased IL-6 levels (Figure S6D), while nuclei maintained large CENP-A foci at centromeres (Figures 5A and 5C). Furthermore, the switch back to basal proportions occurred under conditions in which CIN levels remained unchanged relative to the induced condition (Figure S6E), reinforcing our hypothesis (Figure 2) discarding a prominent role for genome instability in CENP-A-induced EMT. We extended the analysis to the RPE-1 cell line, with only hybrid cells at start, and found that induced mesenchymal states also disappeared in the p53-defective cells after 10 days of reversal (Figure S6F).

**Figure 5.**
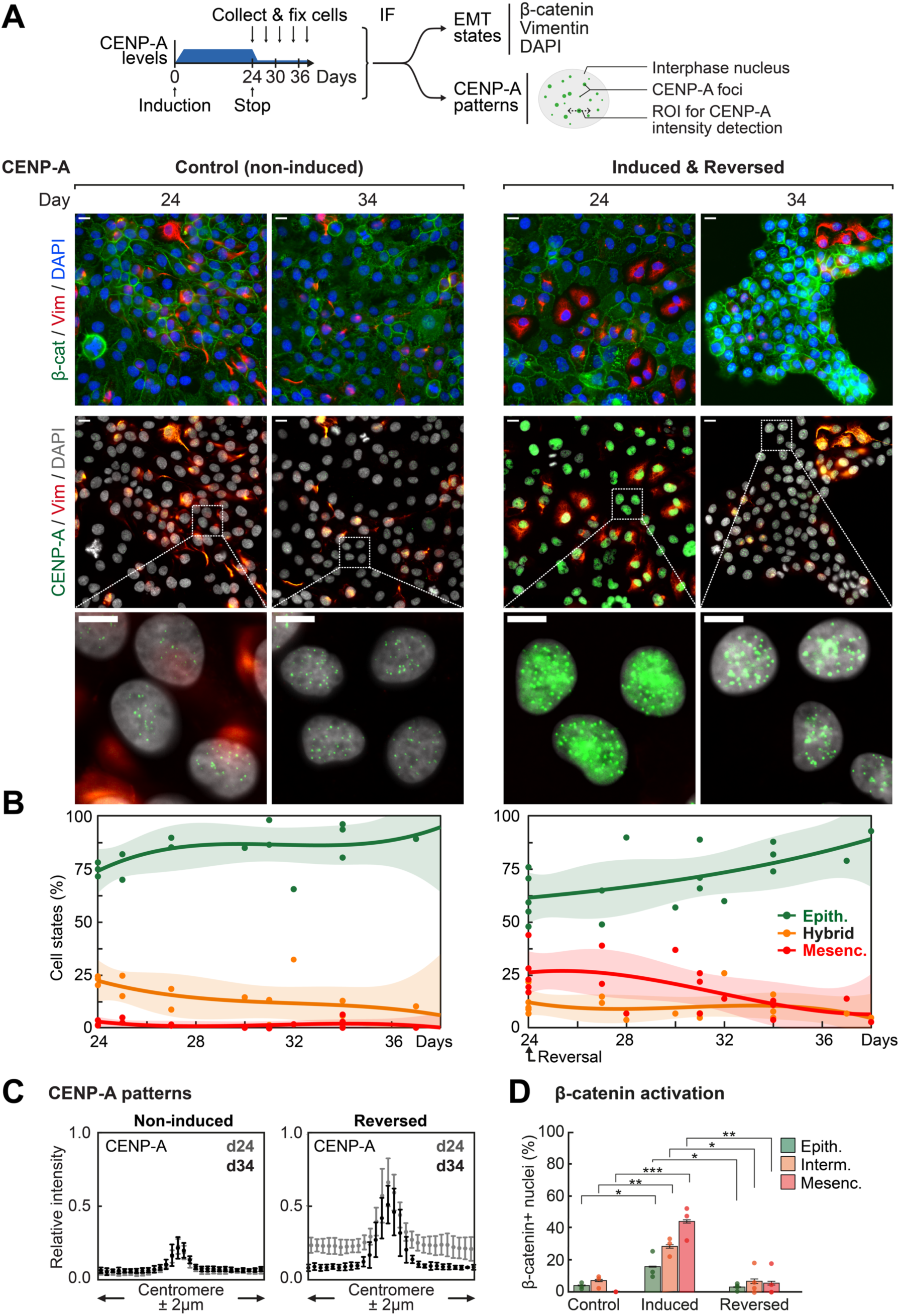
EMT states and subnuclear CENP-A localization following reversion to basal CENP-A levels. (A) *Top:* Experimental strategy. We induced high CENP-A expression (+Dox) until day 24. We then washed and cultivated cells without induction (-Dox) for at least 10 days to allow CENP-A levels to return to baseline. We assessed both EMT states and CENP-A subnuclear localization patterns. EMT quantification for sustained induction is shown in Figure 1 for comparison. ROI: Region Of Interest. *Bottom:* Representative epifluorescence images. Upper panel shows β-catenin (green), vimentin (red), and DAPI (blue). Lower panel shows CENP-A (green), vimentin (red), and DAPI (grey). Scale bars = 20 µm (wide fields), 10 µm (zoomed images). (B) Quantification of epithelial (green), hybrid (orange) and mesenchymal (red) states over time following CENP-A reversal at day 24. Plots show the trendline as polynomial of the third degree ± 95% confidence interval (shaded area) and data points. n > 200 cells counted across ≥ 3 fields for each data point. N = 3 biologically independent experiments. (C) Dot-scan plots displaying the relative intensities of CENP-A IF signal intensity along a 4 μm line traced above representative centromeric chromatin domains (as shown in panel A). (D) Proportions of cells with positive β-catenin nuclear signal at day 34. Plots show mean ± 95% confidence interval for 3 or 4 independent experiments. Dots indicate mean per experiment. n ≥ 600 cells/state across ≥ 3 fields for each replicate. Statistical significance tested by two-tailed Welch’s t test. * = p-value < 0.05, ** = p-value < 0.01, *** = p-value < 0.001. Representative epifluorescence images in Figure S5.

Altogether, these results indicate that the mesenchymal cell population favored upon CENP-A induction result from effects associated to ectopic CENP-A, outside centromeres, rather than its accumulation at centromeres, in line with ectopic CENP-A acting as a reversible epigenetic driver.

## Discussion

While CENP-A has long been considered for its key role in centromere function and genome stability, we unveil here a non-genetic feature of CENP-A in regulating epithelial-mesenchymal plasticity. We summarize our findings in Figure 6. At basal levels, CENP-A specifically marks the centromere and is barely detected ectopically. Under these conditions, the fibrocystic cell population consists in more than 80% epithelial states, 10% hybrid states and less than 5% mesenchymal states. At high CENP-A levels, large CENP-A foci form at centromeres and ectopic CENP-A becomes clearly visible. Under these conditions with ectopic CENP-A, opportunities to enforce EMT occur and mesenchymal states in the population reach 30%. In this context, we identified two distinct cell-cycle dependent EMT trajectories: (**i**) an inflammatory program, pre-existent in control cells, in G1 or outside the cell cycle, and (**ii**) a CENP-A-induced developmental program predominant in S and G2/M phases of the cell cycle. Thus, a window of opportunity for a new EMT arises throughout the cell cycle. Remarkably, interrupting high CENP-A provision erased induced programs along with ectopic CENP-A incorporation, in line with non-genetic alterations. We propose an unconventional, non-centromeric function for CENP-A in epigenetically modulating epithelial-mesenchymal plasticity. Notably, CENP-A removal from chromosome arms leaves cells marked with large CENP-A foci at centromeres. How this mark at centromeres is maintained and impacts cell destinies remains to be evaluated. Nevertheless, these conditions enabled to reverse the balance in the cell population to return to the basal situation, demonstrating a strong dynamic and plastic regulation of EMT programs. Here, we discuss the importance of non-centromeric CENP-A and its relevance for developmental biology and cancer.

**Figure 6.**
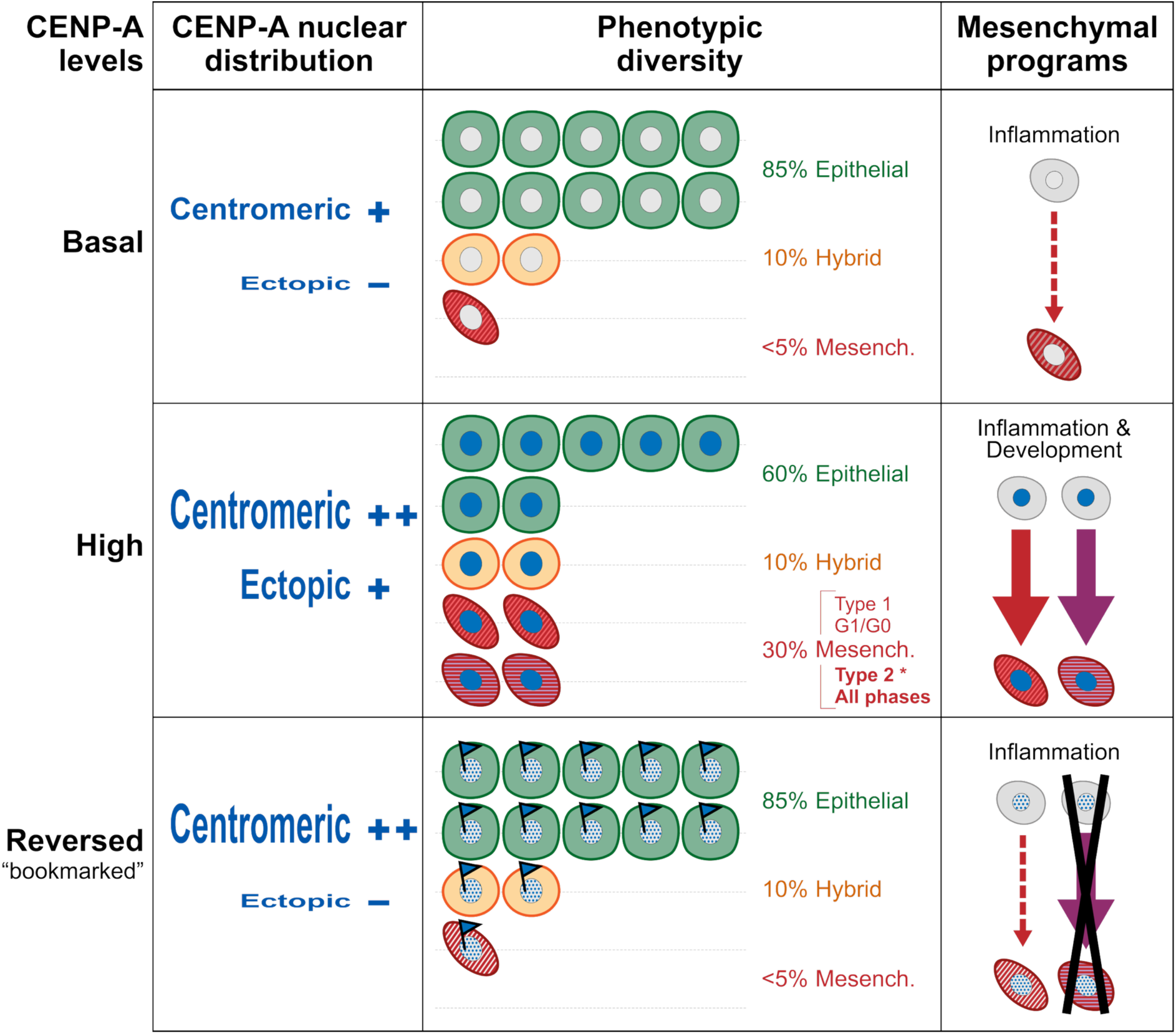
Non-centromeric CENP-A is a reversible epigenetic driver of plasticity. *Left:* CENP-A subnuclear distribution depending on CENP-A levels. At basal levels, only centromeric CENP-A is detected as discrete nuclear foci (small dots) during interphase. Induction of high CENP-A levels leads to increased centromeric CENP-A (big dots) and significant ectopic incorporation at chromosome arms (strong background). Upon reversal to basal expression levels, ectopic CENP-A is lost (no background) while the elevated centromeric levels (big dots) persist for at least 10 days. This marks cells epigenetically, indicating that cells have previously experienced high CENP-A levels. *Middle:* EMT states over time in culture, starting from a mixed population of epithelial, hybrid and mesenchymal states. With basal CENP-A levels, cells favor epithelial states. Elevated CENP-A expression increases the heterogeneity of the population, with various mesenchymal states. Reversal to basal expression levels restores the original distribution of cell states. However, cells retain altered features, including changes in morphology and persistent CENP-A marking. *Right:* EMT trajectories. At basal CENP-A levels, a minority of cells undergo an EMT characterized by an inflammatory gene expression signature. Upon CENP-A induction, hybrid cells can undergo EMT through distinct trajectories, each associated with a unique transcriptomic signature and cell cycle regulation. The inflammatory EMT, associated with G1 or cell cycle exit, is further stimulated/amplified. The developmental-like EMT, associated with all cell cycle phases, emerges exclusively upon CENP-A induction. Reversal to basal expression levels suppresses the developmental-like EMT.

### CENP-A in mediating phenotypic plasticity and heterogeneity

This unanticipated role for CENP-A as a regulator of cell plasticity to promote EMT raises a central question: *how does ectopic CENP-A drive this phenotypic plasticity?* Since this capacity of CENP-A to drive EMT only operates when hybrid states are detectable before induction (Figures 1B, 1C and S1H), we propose that CENP-A enables (or “unlocks”) transitions toward mesenchymal states in cell populations harboring hybrid states. Thus, hybrid cells, poised to generate phenotypically diverse progeny^45,72–74^, could represent a pool of cells responsive to high CENP-A levels. It is possible that the chromatin replication-dependent error correction mechanism which normally removes ectopic CENP-A^17^ becomes saturated, thereby enabling stable accumulation of CENP-A at sites poised to promote EMT. Furthermore, our data discard an evolution of cell populations driven solely by genetics. Indeed, the emerging mesenchymal population showed modest increase in CIN which could not account for the observed phenotypic shift (Figure 2B). In control conditions, our fibrocystic population predominantly exhibited epithelial and hybrid states, rare cells completing EMT along a single trajectory coupled to cell cycle exit (Figure 4B-E). Consistently, whereas EMT cells exist in S/G2/M phases during mouse neural crest development^41^, EMT has been tied to cell cycle exit in fibrosis with cells typically remaining in hybrid states^39,40^. However, we discovered that CENP-A enables cells to follow at least two distinct trajectories, coupled to specific cell cycle stages, toward mesenchymal states (Figures 3A and 4C-F). This aligns with the view of EMT as a context-dependent spectrum of cell states^31,75^. Here, the flexible chromatin landscape provided by CENP-A incorporation may enable stochastic exploration of alternative fates^76^. This could exploit the possibility to exit at any time during the cell cycle to change destinies. Indeed, the fact that we discovered new directional dynamics following CENP-A induction introduces an additional layer of plasticity, with an expanded range of possible trajectories for cells. To our surprise, while many genetic defects could have occurred, and despite persistent CIN and elevated centromeric CENP-A, restoring CENP-A expression to basal levels can restore the initial phenotypic distribution in the cell population (Figure 5A-B). These findings further argue against a model in which permanent genetic alterations drive EMT, instead pointing to a reversible, non-genetic mechanism mediated by ectopic CENP-A deposition, as the reversion occurred concomitantly with the loss of ectopic CENP-A (Figure 5C). Considering that excess CENP-A, through unconventional association with H3.3 chaperones, allows the formation of heterotypic CENP-A/H3.3 nucleosomes at genomic regions with high histone turnover^21^, we examined if we had increased CENP-A at critical regulatory locations. Our ChIP data showed significant CENP-A gains at EMT genes preceding the EMT and correlating with transcriptional changes (Figure 3A, D). We thus propose that ectopic CENP-A deposition provides an enhanced chromatin plasticity enabling expression programs that govern epithelial-mesenchymal plasticity. The very structure of the ectopic particle, partially opened^77^, could make chromatin more accessible, and thus corresponding regions more fluid, thereby conferring a higher probably to switch to alternative states. This plasticity, here driven by CENP-A modulation, extends broadly the view of chromatin as a versatile landscape^78^ in which histone chaperones, by acting on H3 variants deposition^79^, or the balance in the proportions of chromatin-bound H3.1 and H3.3^80^ can impact cell fate. Changing histone dosage could similarly affect cell destinies^81^. This global modulation is reminiscent of concepts of epigenetic noise^82,83^ and epigenetic regulatory network interferences^84^, and enriches prevailing deterministic models of EMT. In this framework, excess CENP-A may saturate the replication-coupled removal of ectopic CENP-A, thereby increasing epigenetic noise.

### Relevance in development and cancer

To date, the elevated expression of CENP-A in tumor cells has largely been attributed to its role in cell proliferation and CIN. The finding that CENP-A not only amplifies a pre-existing inflammatory program in fibrocystic MCF10-2A cells^39,40,47^ but also induces a distinct developmental-like program (Figures 3A and 4C-F) was a real surprise. These novel properties may be related to new options offered by cell cycle dynamics and the possibility to engage into EMT at different stages. These findings gain significance when considering new roles of CENP-A in promoting cellular plasticity, as it may involve critical time windows for cell fate changes, such as during early development and cancer progression. Thus, our data provide a framework to explain unexpected, chromosome segregation-independent functions of CENP-A. In this context, it will be exciting to revisit the *de novo* assembly during the differentiation of non-dividing *Drosophila* midgut epithelial cells^85^ and its retention in post-mitotic neurons^86^. Likewise, both embryonic^87^ and induced pluripotent^88^ stem cells require high CENP-A mRNA levels to differentiate, whereas CENP-A depletion in cardiac progenitors induces apoptosis upon lineage commitment^89^. In the context of tumorigenesis, our findings also recapitulate key features of tumor progression^90–93^, including the extensive intratumoral heterogeneity reported in glioblastoma^94^, where tumor-initiating cells exhibit elevated CENP-A levels^95^. EMT heterogeneity correlates with metastatic potential in mouse models and patient tumors^96–98^. Current models propose that metastatic competence relies on the synergy between multiple EMT states and programs to confer increased adaptability^99–101^. These features may explain the strong correlation of high CENP-A levels with cancer aggressiveness and metastatic progression (Table 3). Importantly, the reversibility that we found for CENP-A (Figure 5B) is in contrast with recent observation of transient depletion of Polycomb proteins^102^, yet it underlines another important aspect of non-genetic driven mechanisms of importance in tumorigenesis. In addition, it indicates that stable ectopic CENP-A incorporation drives this phenotypic plasticity. This is particularly relevant since non-centromeric CENP-A deposition is likely a widespread phenomenon in cancer, driven not only by elevated expression (∼50% of cancers^21^) but also by dysregulation of H3 variant network regulators such as CHAF1B^25^, DNAJC9^26,27^ and NuRD^103^. Furthermore, while metastatic tissues can lose the high CENP-A expression signature (cBioPortal datasets), we found that CENP-A subnuclear staining enables to identify cells that had previously experienced elevated CENP-A levels, even several days after reversal (Figure 5A-C). This CENP-A accumulation at centromeres could influence centromere function^104^, boundary definition and stability across cell divisions^105,106^. Future studies leveraging recent technologies such as DiMeLo-seq^107^ will be critical for high resolution mapping. Moreover, this marking could serve as a marker of prior CENP-A upregulation, providing insights into the temporal dynamics of CENP-A regulation throughout tumor evolution. These persistent marks might also serve as biomarkers for metastatic tumors. One could envisage a scheme in which transient CENP-A upregulation enables cell migration, followed by a return to basal levels facilitating tumor development at a distant site in a favorable environment. Therefore, erasing ectopic CENP-A by interfering with the chaperones DAXX and/or HIRA represent potential approaches to prevent tumor progression and dissemination. This could take advantage of recent technologies such as proteolysis-targeting chimeras (PROTACs) and the development of targeted inhibitors.

**Table 3.** Correlation between high CENP-A levels and cancer aggressiveness.

| **Cancer type** | **Correlation** | **References** |
| --- | --- | --- |
| Breast | Grade, **invasion**, **recurrence**, poor survival, **metastasis** | 111–116 |
| CNS | Grade, **recurrence**, poor survival, **immune evasion** | 115,117–119 |
| Ovarian | Grade, poor survival, **immune evasion** | 116,120,121 |
| Lung | Grade, **recurrence**, poor survival, proliferation | 115,116,122,123 |
| Liver | Grade, proliferation, size, **invasion** | 124,125 |
| Gastric | Poor survival, **metastasis** | 115,116,126 |
| Osteo-sarcoma | Size, poor survival, proliferation, **metastasis** | 127 |
| Prostate | Grade, poor survival, **metastasis** | 128,129 |
| Kidney | Grade, poor prognosis, **metastasis** | 130 |
| Head & Neck | **Metastasis** | 131 |

In conclusion, our work unveils a new non-centromeric function for CENP-A in promoting epithelial-mesenchymal plasticity by enabling transition between alternative cell states in a reversible way. While we focused on local effects targeting transcriptional regulation, non-centromeric CENP-A may also affect, either directly or *via* long-range effects, replication, higher-order structures, and stemness properties, all modulated by histone variants equilibrium^108–110^. These findings open exciting avenues to integrate the importance of non-centromeric CENP-A in cell fate decisions during development and cancer progression.

## Methods

### Cell lines

All parental human cell lines were originally obtained from ATCC: MCF10-2A (CRL-10781), HCC1954 (CRL-2338), MCF7 (HTB-22), T47D (HTB-133) and hTERT RPE-1 (CRL-4000, kind gift from Daniele Fachinetti, Institut Curie). The MCF10-2A cell line was authenticated by STR profiling (Powerplex 16 HS). All cell lines were transduced with the inducible CENP-A overexpression construct Tet-On–CENPA–FLAG–HA as in Jeffery et al.^19^. All cell lines were tested negative for mycoplasma contamination. For generation of p53-WT and p53-DN cells, we transduced cells with an empty vector (pWZL Hygro; Scott Lowe, Addgene plasmid #18750) or a vector containing a dominant-negative TP53 construct (pBABE-Hygro p53DD, Bob Weinberg, Addgene plasmid #9058) as in Jeffery et al.^19^.

### Cell culture

We cultured all cells at 37°C in 5% CO_2_. MCF10-2A cells were cultured with DMEM:Ham’s F12 5% horse serum, 5 ng/ml epidermal growth factor, 100 ng/ml cholera toxin, 0.01 mg/ml insulin, 500 ng/ml hydrocortisone and 1% Pen/Strep; HCC1954 and T47D cells with RPMI Medium 10% FCS and 1% Pen/Strep; MCF7 cells with DMEM 10% FCS and Pen/Strep 1%; RPE-1 with DMEM:Ham’s F12 10% FCS, 0.123% sodium bicarbonate and 1% Pen/Strep. All base media were purchased from ThermoFisher Scientific.

### Induction of high CENP-A expression

We induced high CENP-A expression by adding doxycycline (Dox; dissolved in DMSO) to growth media. A concentration of 10 ng/ml was defined as 1X and used in all experiments unless otherwise specified (10X Dox). Control cells received DMSO alone. Growth media (±Dox) was refreshed every 2-3 days. To restore basal CENP-A expression, Dox-containing medium was removed, cells were washed twice with pre-warmed PBS and cultured in Dox-free medium.

### Cell extract preparation and western blotting

We performed Western bloting as described in Jeffery et al.^19^ with protein extracts from ∼30 000 cells (considered 1X load) per gel lane. **Primary antibodies:** active Beta-catenin (D13A1) 1:100 (#ab16051 Abcam), CENP-A 1:500 (#2186 Cell Signaling Technology), human IL-6 1:300 (#AF-206-NA, R&D Systems), Vimentin (R28) 1:1000 (#3932 Cell Signaling Technology), Vinculin 1:10000 (#V9131 Sigma-Aldrich), yH2A.X 1:1000 (# 05-636 Merck). **Secondary antibodies:** Jackson ImmunoResearch, 1:10000, donkey anti-mouse, donkey anti-rabbit, donkey anti-goat.

### Immunofluorescence microscopy

We grew cells on glass coverslips pre-coated with collagen + fibronectin (each at 1 µg/ml) in culture dishes. For cell division tracing, cells were pulse-labeled cells with 5 µM CFSE according to the manufacturer’s instructions. Cells were fixed with 2% PFA, permeabilized with 0.2% Triton X-100 in PBS for 5 min, blocked with 5% BSA plus 0.1% Tween-20 in PBS, and incubated with primary and secondary antibodies, followed by DAPI counterstaining. For CENP-A and CENP-B co-staining (Figure S1A), we performed Triton-CSK extraction prior to fixation. We mounted coverslips in Vectashield medium and used a Zeiss Axiovert Z1 microscope with CoolSNAP HQ2 camera, with MetaMorph software. For image-based data visualization, same intensity (pixel ranges) was applied across conditions, except for Figure S1B, 5A and S6A, where CENP-A staining in non-induced cells was displayed with a greater apparent intensity to detect CENP-A foci, and in Figure S1H, where vimentin in the MCF10-2A cell line was displayed with a reduced apparent intensity relative to other conditions. **Primary antibodies:** Beta-catenin 1:100 (#9562 Cell Signaling), CENP-A (3-19) 1:300 (#ADI-KAM-CC006-E Enzo Life Sciences), cleaved Caspase-3 1:400 (#9661 Cell Signaling), E-cadherin (24E10) 1:200 (#3195S CST), Vimentin (N-term) 1:200 (#5741S Progen). **Secondary antibodies:** 1:1000 Alexa Fluor donkey anti-mouse IgG (H + L) 488 or 647, 1:1000 Alexa Fluor goat anti-rabbit IgG (H + L) 488 or 594, 1:1000 Alexa Fluor goat anti-Guinea Pig IgG (H + L) 594.

### Doubling time estimation

CFSE fluorescence intensity decreases by half with each cell division (i.e., cells in generation 1 display approximately 50% of the initial signal, cells in generation 2 approximately 25%, etc.) Based on CFSE signal intensities, we determined the proportion of cells in each generation. For instance, after 96h in culture, cells with CFSE intensity ∼0.25 (generation 2) have undergone two divisions, thus yielding an estimated doubling time of 96/2 = 48h. Considering the fractions of cells in each generation, we calculated the average doubling time for the entire cell state population.

### Quantification from microscopy images

Images were acquired from at least three fields per coverslip using the DAPI channel. Between 200 and 1000 nuclei were counted per time point and condition and EMT states were determined based on epithelial and mesenchymal marker staining, using ImageJ software. For vimentin intensity quantification in hybrid cells, we generated masks from at least three fields for the red (vimentin) and green (E-cadherin or β-catenin) channels and combined them using the “AND” command to identify regions of co-expression. From the red channel, we extracted the integrated density (IntDen; sum of all pixel values /area size) and normalized to the number of hybrid cells. Micronuclei were quantified in ImageJ from at least three fields, with more than 1000 nuclei analyzed per condition.

### FACS analysis

For S phase-labelling, we pulse-labeled cells in culture with 10 µM 5-ethynyl-2′-deoxyuridine (EdU) for 20 min, as in Mendiratta et al.^132^. For all analyses, we counted cells using a Vi-Cell XR Cell Viability Analyzer (Beckman coulter). 1 million cells were harvested by trypsinization and fixed with 70% cold ethanol. After washing with 1 ml PBS, cells were blocked in 5% BSA plus 0.1% Tween-20 in PBS for 45 min at room temperature. We then added primary antibodies (1 µl each): E-cadherin (24E10) (#3195S CST), Vimentin (VI-RE/1) conjugated with APC (#MA5-28601 ThermoFisher), or isotype control (IgG1κ conjugated with APC (#17-4714-42, ThermoFisher). After 1 h, secondary antibody (1 µl Alexa Fluor 488-conjugated goat anti-rabbit IgG (H+L)) was added for E-cadherin detection and incubated for an additional 1 h (room temperature, protected from light). Cells were washed twice in 1 ml PBS-Tween and stained with 0.5 µg/ml DAPI. For EdU detection, we used the Click-iT EdU Cell Proliferation Kit with Alexa Fluor 488 dye (#C10337, ThermoFisher) according to the manufacturer’s instructions, prior to DAPI staining. For EdU-labeled cells, we acquired data using an Aria III cytometer (BD Biosciences) at the Cytometry platform CYTPIC of Institut Curie. Otherwise, we acquired data with an Attune NxT cytometer (ThermoFisher Scientific). We performed analysis on a minimum 15,000 gated cells, excluding cell debris and doublets, with FlowJo software (v10.9.0).

### Bulk RNA-seq analysis

Bulk RNA sequencing data were obtained from Jeffery et al. (2021)^19^. We aligned RNA-seq reads to the human reference genome (GRCh38) using HISAT2 (v2.2.0 beta) in paired-end mode with default parameters and Ensembl gene annotations (release 104). We computed gene-level counts from primary alignments with MAPQ > 2 using featureCounts (Subread v2.0.2). To identify differentially expressed genes, we used DESeq2 (v1.46.0) on raw counts, considering batch effects, p53 status, CENP-A induction, and the synergistic effect of p53 status and CENP-A induction, relative to control (p53-WT and no induction). Genes were defined as differentially expressed at an adjusted p-value < 0.05. All analyses were performed using custom R scripts with DESeq2, rtracklayer, and tidyverse packages. For the second, biologically independent RNA-seq analysis in Figure S3A, pseudo-bulk gene expression was estimated from the RNA modality of our multi-omics dataset. We aggregated counts across cells from the same condition, normalized by library size, and applied log-transformation.

### Bulk ChIP sequencing

We performed chromatin immunoprecipitation sequencing (ChIP-seq) for CENP-A and H3.3 using the native nucleosome isolation procedure described in Gatto et al.^108^ with minor modifications. We grew p53-DN MCF10-2A TetOn-CENPA-FLAG-HA cells with 0X Dox (DMSO control), or with 1X Dox (10 ng/ml) for 24 days. We harvested samples at days 10 and 24. All steps were performed at 4°C with Protease inhibitors (Roche) included in all buffers to prevent proteolytic degradation. For each IP, 2-5 million cells were harvested and processed in aliquots of 1 million cells per tube by resuspending cell pellets in 100 µL ice-cold lysis buffer (50 mM Tris–HCl, pH 7.5; 150 mM NaCl; 5 mM CaCl₂; 1% Triton X-100; 0.5% NP-40; and EDTA-free protease inhibitors, Roche) for a 4-min incubation at room temperature. For each IP reaction, we prepared antibody-coupled Protein A Dynabeads (100 µL per IP; Invitrogen, 10002D) by blocking for 3h at room temperature in 1 mL blocking buffer (PBS with 0.1% Tween-20, 2.5% filtered BSA and 0.2 µg tRNA), with gentle rotation, washing once with PBS-T (0.02% Tween-20) and incubating for 1h at room temperature in 200 µL PBS-T with 5 µg anti-H3.3 (Active Motif, 91191), 10 µg anti-CENP-A (Cell Signaling, 2186), or 10 µg control IgG (Abcam, ab46540). After a wash in binding buffer, we added the antibody-coupled beads to the chromatin and incubated overnight at 4°C on a rotating wheel. The next day, beads were recovered using a magnetic rack and sequentially washed twice with Wash Buffer 1 (10 mM Tris-HCl, pH 8.0; 140 mM NaCl; 1% Triton X-100; 0.5% NP-40; 0.1% SDS), twice with Wash Buffer 2 (as above but with 360 mM NaCl), twice with Wash Buffer 3 (as above but with 250 mM LiCl and 0.5% Triton X-100), and twice with TE (10 mM Tris–HCl, pH 8.0; 1 mM EDTA). Beads were resuspended in 20 µL TE and treated with RNase A (2 µL, 10 mg/mL, 30 min at 37°C) and then proteinase K (2 µL, 20 mg/mL) in the presence of 2.5 µL 2% SDS (1 h at 37°C). DNA was purified using AMPure XP beads (Beckman Coulter, A63880) as per the manufacturer’s protocol and eluted in 20 µL nuclease-free water. Quality and concentration were assessed using a TapeStation 4200 (Agilent) and a Qubit 3.0 fluorometer (Invitrogen). Sequencing libraries were then prepared with 15 ng DNA using the Illumina TruSeq ChIP kit and sequenced on the Illumina NovaSeq 6000 (PE100) at the Institut Curie Next Generation Sequencing (NGS) platform.

### ChIP-seq analysis

We aligned reads to the GRCh38 reference with Bowtie2 (v2.4.4, paired-end mode, very-sensitive preset). We flagged PCR duplicates using SAMtools (v1.19.2) after sorting then we indexed the corresponding BAM files. We computed fragment coverage using the genomecov function from BEDtools (v2.30.0). We summed coverage over 5 kb windows centered at transcription start sites (TSSs), normalized the values to the total number of mapped reads (counts per million), and normalized to input. We applied log10 transformation followed by z-score normalization (relative to the mean signal and standard deviation). For genome-wide distribution analysis, we computed the average signal in consecutive, non-overlapping 100 kb bins. We performed downstream analyses in R using custom scripts and the packages rtracklayer (v1.66.0) and tidyverse (v2.0.0). For hierarchical clustering analysis, we averaged signals at TSSs across non-induced (control) samples to reduce noise. For each timepoint, we then computed the difference (induced - non-induced) for each gene and clustered genes by the relative gain/loss of CENP-A and H3.3 at their TSS. We used hierarchical clustering analysis via “complete” aggregation method applied on Euclidian distance. We defined cluster boundaries by cutting the dendrogram at the height yielding eight distinct clusters. We assessed enrichment for Gene Ontology (GO) Biological Process terms using the enrichGO function from the clusterProfiler R package (v4.14.6) with annotations from the org.Hs.eg.db database. We considered terms with p < 0.05 as significantly enriched. We visualized the results using the plotEnrich function from the enrichR R package (v3.4).

### Single-nucleus multiomics

We grew p53-DN MCF10-2A TetOn-CENPA-FLAG-HA cells with 0X Dox (DMSO control), or with 1X Dox (10 ng/ml) for 24 days. We harvested samples at days 10 and 24. We prepared samples according to the Active Motif Single-Cell Multiome Sample Preparation Protocol. In brief, we harvested cells by trypsinization, resuspended in complete media and mixed thoroughly by pipette, centrifuged at 500g at 4°C to pellet the cells and remove supernatant, resuspended cells in the appropriate volume of ice-cold cryopreservation solution (50% horse serum, 40% growth media, 10% DMSO) to achieve a concentration of 4 million cells/mL, then transferred 500 uL (2 million cells) to a 1.5 mL Eppendorf tube on ice and froze cells by transferring the tubes to a pre-chilled container and placing at-80°C. The samples were then shipped in dry ice to Active Motif services, who proceeded to a cell viability test by Trypan blue and nuclei isolation with 5 minutes lysis as described in the Nuclei Isolation for Single Cell Multiome protocol. Nuclei concentration and quality were confirmed with PI and Acridine orange staining. ATAC and RNA libraries were generated using the Chromium Next GEM Single Cell Multiome ATAC + Gene Expression protocol from 10X Genomics, relying on microdroplet-based isolation of single nuclei.

### Single-nucleus multiomics analysis

**Data processing:** We aligned reads using CellRanger ARC (v2.0.2) to the GRCh38 reference (index file provided by 10X Genomics, v2020-A-2.0.0). For the snRNA modality, we processed gene expression matrices using Seurat. We filtered out low-quality cells and retained cells with ≥ 200 features using specific thresholds per sample, based on the number of UMIs (3.5-40k), detected features (2-10k) and mitochondrial DNA percentage (4-30%). We normalized read counts with LogNormalize (scale factor 10000) and assigned cell cycle phases using CellCycleScoring. We identified highly variable genes using FindVariableFeatures and performed principal component analysis (PCA), retaining the top 50 components. For the snATAC modality, we processed ATAC fragments using ArchR (v1.0.2). We created Arrow files (≥ 4 TSSs and ≥ 1000 fragments detected). We computed Tile and GeneScore matrices, removed doublets using addDoubletScore (k = 10) and filterDoublets, and filtered cells by fragment count (min 5.6; max 63k). We computed latent semantic indexing (LSI) using addIterativeLSI and identified marker peaks with getMarkerFeatures (Wilcoxon test; covariates: TSS enrichment, log10 fragments), applying FDR ≤ 0.01 and log₂FC ≥ 1. **Multi-omics data integration:** We performed paired data integration and visualization with Multi-Omics Wasserstein inteGrative anaLysIs (Mowgli, v0.3.1; github.com/cantinilab/Mowgli)^65^. Mowgli combines Non-negative Matrix Factorization (intNMF) and Optimal Transport (OT) to integrate snRNA and snATAC data. We selected Mowgli due to the demonstrated superior performance of intNMF in clustering multi-omics data^63^, and because OT has been shown to better capture similarities between single-cell omics profiles^64^. This strategy enhances both the performance and the biological interpretability of the results. We used the set of 15614 high-quality cells after single-cell data preprocessing. snRNA gene counts were log normalized using scanpy^133^ version 1.9.6. snATAC peaks were log normalized with TF-IDF method from the muon package^134^ 0.1.5. For snRNA, we used the set of 2589 highly variable genes identified in previous analyses. For snATAC, we selected the top 10000 highly variable peaks associated with the promoter regions and the top 10000 highly variable peaks related to distal regions. We applied Mowgli^65^ version 0.3.1 to the three obtained matrices: snRNA and snATAC on promoter regions and snATAC on distal regions. Standard parameters were used, and the number of latent dimensions was set to 100 to increase the granularity of the integration. In the following analyses, we used the multi-omics cell embeddings outputted by Mowgli. We performed Leiden clustering^135^ at resolution 1.1 with the scanpy Leiden wrapper. Distances across clusters were computed using scanpy dendrogram method with default parameters. Differentially expressed genes were assessed using the scanpy rank_genes_groups method, using a wilcoxon test to assess the statistical significance and a Benjamini-Hochberg correction to adjust p-values. Cell scores for epithelial and mesenchymal states and cell cycle were measured using the score_genes method. **RNA velocity:** Spliced and unspliced counts from the snRNA modality were measured with velocyto^136^ version 0.17.17 using the run_10x wrapper. The loom files outputted by velocyto were then used to measure velocity with scVelo^66^ using the dynamical model. Finally, velocity vectors were summarized with CellRank^137^ version 2.0.4 using the ConnectivityKernel method. We used the neighboring graph derived from the top 10 nearest neighbors per cell in the multi-omics cell embeddings of Mowgli by giving it a weight of 0.2 when combining it with the velocity kernel (that was given a weight of 0.8). We plotted the velocity field on the multi-omics cell embeddings previously outputted by Mowgli.

### Gene regulatory network inference and TF activity analysis

We performed gene regulatory network (GRN) inference from the Multiome dataset using HeterogeneoUs Multilayers for MUlti-omics Single-cell data (HuMMuS, v0.1.7; github.com/cantinilab/HuMMuS)^70^. HuMMuS leverages heterogeneous multilayer networks to reconstruct regulatory mechanisms by integrating both inter-and intra-omics interactions. First, we inferred an RNA-based GRN with GRNBoost2^138^ using the wrapped version stored in SCENIC^139^ (v0.12.1) and a peak-peak network from single-cell ATAC peaks with Circe (v0.3.6; github.com/cantinilab/circe), a scalable tool for cis-regulatory inference with performances close to those from Cicero^140^. We used the cell embedding derived from Mowgli to aggregate the metacells. For HuMMuS, we selected the top 50 000 TF-target links from GRNBoost2 and the top 300 000 peak-peak interactions from Circe according to their score, and employed the internal wrapper to compute a TF-TF interaction layer using Omnipath^141^. From the resulting multilayer GRN, we estimated TF activity with the univariate linear model implemented in decoupler^142^ (v2.0.4) and computed differential TF activity between clusters 7 and 12 using the rankby_group method implemented in decoupler. The top 10 most differentially active TFs (adjusted p ≤ 0.05) in each cluster were retained for downstream analyses. For each cluster, we built a cooperativity network across the top TFs using their 200 highest-scoring targets (as ranked by HuMMuS).

## Acknowledgments

We thank Dr. Jean-Pierre Quivy and Dr. Daniele Fachinetti at Institut Curie for their useful critical comments. We thank Dr. Daniel Jeffery and our team for discussions. We thank the PICT-IBiSA@Pasteur Imaging Facility (member of the France Bioimaging National Infrastructure, ANR-24-INBS-0005 FBI BIOGEN), the NGS platform ICGex, and the Cytometry platform CYTPIC of Institut Curie. The work of G.A. was supported by the ERC-2015-ADG 694694 “ChromADICT”, La Ligue Nationale contre le Cancer (labellisation), government grants managed by the Agence Nationale de la Recherche (“Investissements d’Avenir” program, ANR-11LABX-0044 DEEP, ANR-10-IDEX-0001-02 PSL; “France 2030” program, ANR-24-EXCI-0001, ANR-24-EXCI-0002, ANR-24-EXCI-0003, ANR-24-EXCI-0004, ANR-24-EXCI-0005), and by the SiRIC-Curie program “INCa-DGOS-Inserm-ITMO Cancer_18000”. The work of L.C. was founded by the ERC StG MULTIview-CELL 101115618 and a government grant managed by the Agence Nationale de la Recherche (“Investissements d’Avenir” program, ANR19-P3IA-0001, PRAIRIE 3IA Institute). C.R.-P. was funded by Sorbonne Université and by the Fondation pour la Recherche Médicale (Grant No. FDT202304016871).

## Author contributions

G.A. and C.R.-P. conceived the overall strategy, designed the approaches and wrote the manuscript.

G.A supervised the work. C.R.-P. performed cell biology, imaging and biochemistry experiments, and analyzed the corresponding data. A.F. and C.R.-P. carried out cytometry and ChIP assays. S.L. and C.R.-P. analyzed bulk RNA and ChIP-seq data. D.C. and L.C. conducted multiomic data integration and analyzed the results together with C.R.-P., S.L., and G.A. All authors contributed to critical reading and data discussion.

## Supplemental Figures

**Figure S1.**
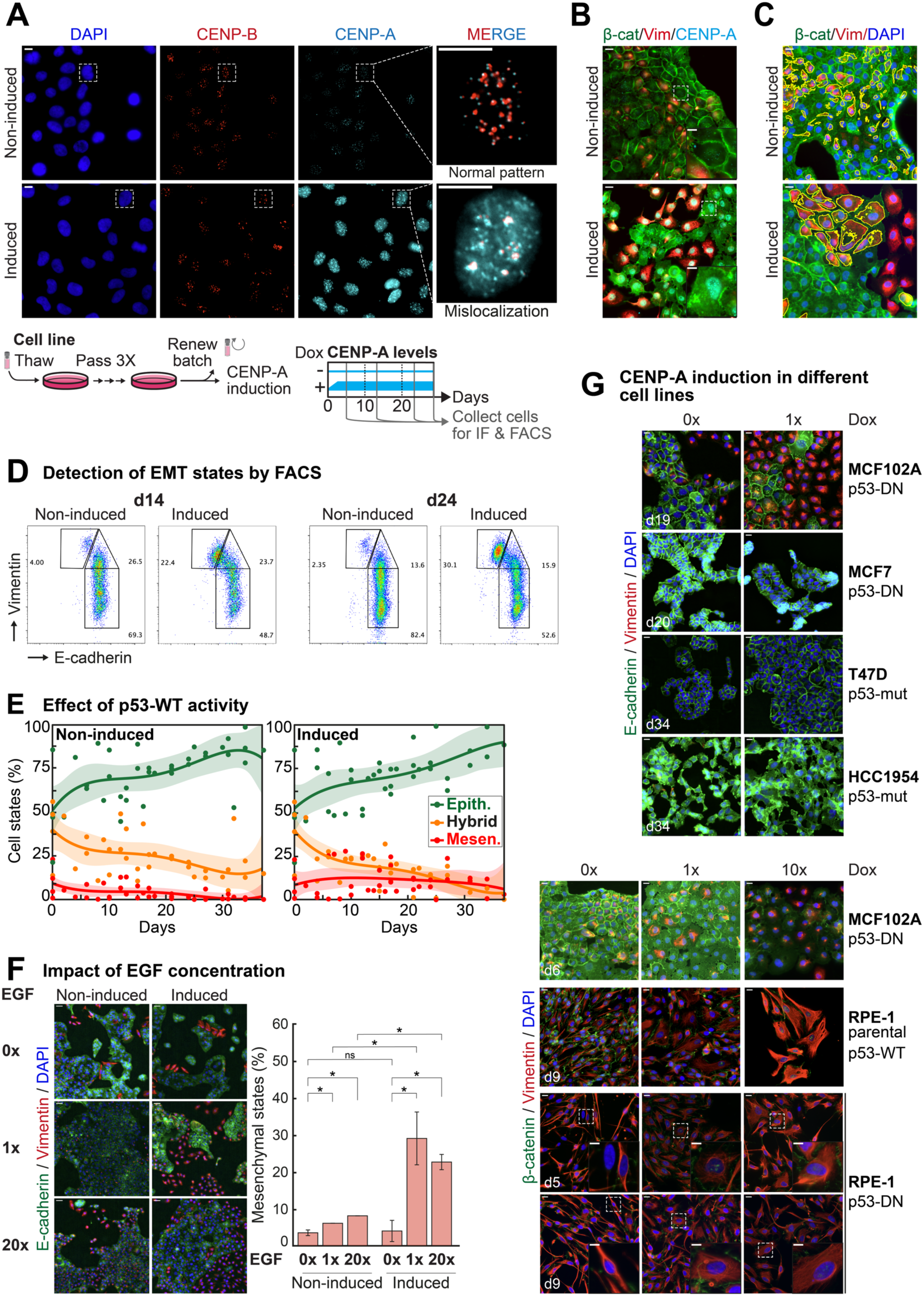
Influence of p53 status, EGF levels and cell type on CENP-A-induced EMT. (A) *Top:* CENP-A localization patterns in interphase nuclei after CSK-Triton extraction. *Left:* Representative epifluorescence images of cells immunostained for CENP-B (red) and CENP-A (cyan), with DNA counterstained with DAPI (blue). Scale bars = 20 µm. Squared boxes indicate cells on the right. *Right:* Insets of representative nuclei. Scale bars = 10 µm. *Bottom:* Experimental strategy (related to Figure 1A). Cells were thawed from frozen stocks and passaged three times before preparing fresh stocks, kept at-80 °C. The rest of the cells was used for the induction. Samples were collected at various time points for IF or FACS analysis. (B) EMT states and corresponding CENP-A patterns. Epifluorescence images (related to Figure 1C) showing β-catenin (green), vimentin (red), and CENP-A (cyan). Insets show representative CENP-A patterns. Scale bars = 20 µm (wide fields), 10 µm (insets). (C) Hybrid EMT states. Epifluorescence images (related to Figure 1D) showing β-catenin (green), vimentin (red), and DAPI (blue). Vimentin staining in hybrid cells (co-expressing β-catenin and vimentin) is segmented (yellow outlines). This enables to monitor average vimentin intensity in hybrid cells across different fields. Scale bars = 20 µm. (D) FACS profiles at days 14 and 24 showing epithelial (E); hybrid (H) and mesenchymal (M) states based on vimentin and E-cadherin expression levels. (E) EMT states across time in p53-WT MCF10-2A cells. Proportions of epithelial (green), hybrid (orange) and mesenchymal (red) states. Plots show the trendline as polynomial of the fourth degree ± 95% confidence interval (shaded area). n > 200 cells across ≥ 3 fields for each data point. N ≥ 5 biologically independent experiments. (F) EMT states 20 days post-induction under different EGF concentrations. *Left:* Representative epifluorescence images showing E-cadherin (green), Vimentin (red) and DAPI (blue). Scale bars = 100 µm. *Right:* Proportions of mesenchymal cells. Plots show mean ± 95% confidence interval. N = 2 biologically independent experiments. Statistical significance tested by two-tailed Welch’s t-test. * = p-value < 0.05. (G) Impact of increased CENP-A levels in different p53-defective cell lines. *Top:* Cells after chronic CENP-A overexpression compared with MCF10-2A cells at day 19. Epifluorescence images show E-cadherin (green), vimentin (red), and DAPI (blue). Scale bars = 20 µm. DN: dominant-negative p53 vector; mut: p53 loss-of-function mutation. *Bottom:* Cells within 10 days of induction, stained for β-catenin (green), vimentin (red), and DAPI (blue). Squared boxes indicate cells on the bottom (insets). 1X Dox = 10 ng/ml, 10X Dox = 100 ng/ml. Scale bars = 20 µm (wide fields), 10 µm (insets).

**Figure S2.**
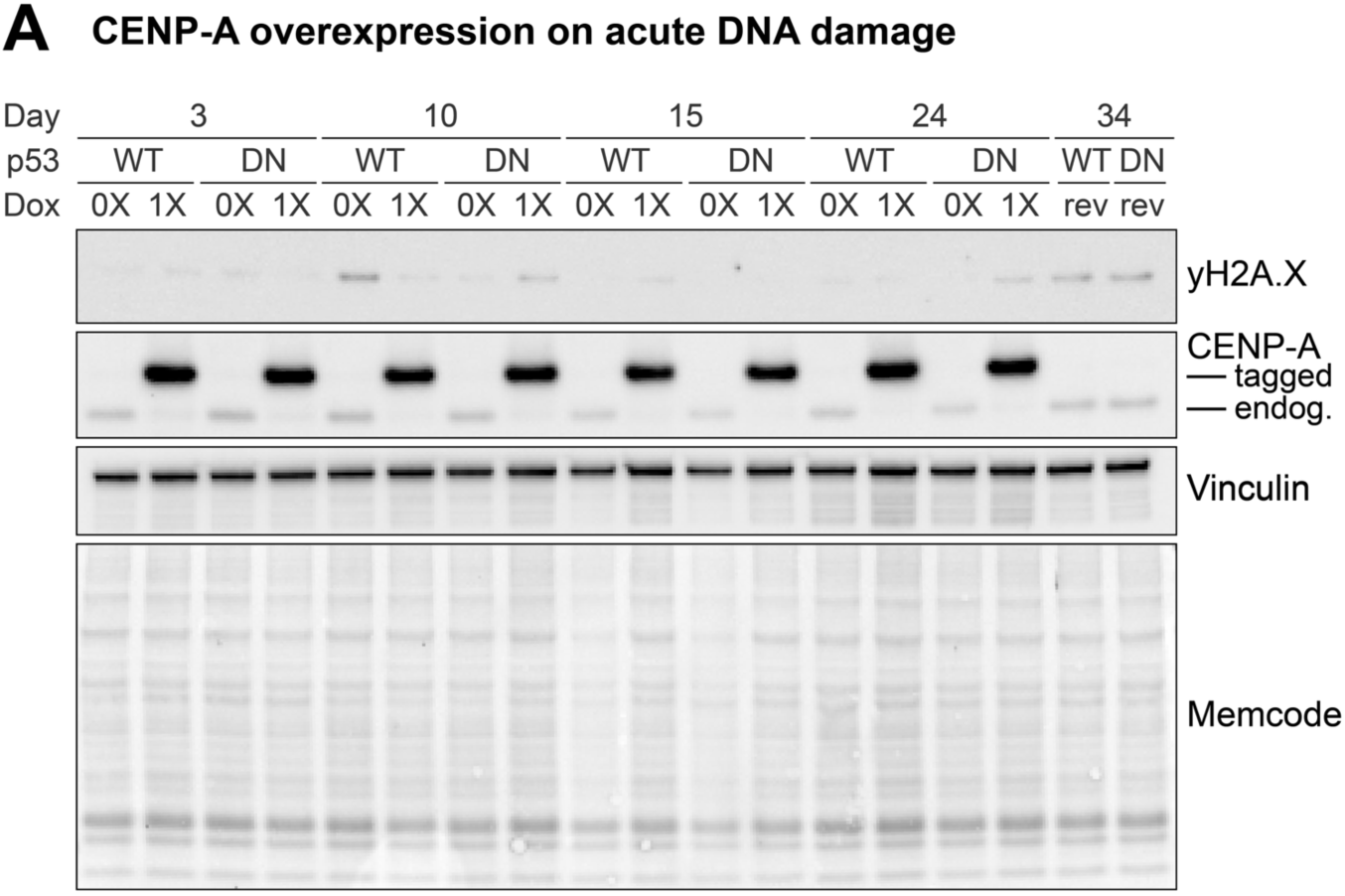
Impact of high CENP-A levels on acute DNA damage. (A) Western blot analysis of total protein extracts from MCF10-2A cells (-Dox or +Dox), with vinculin as loading control. Primary antibodies are indicated on the right.

**Figure S3.**
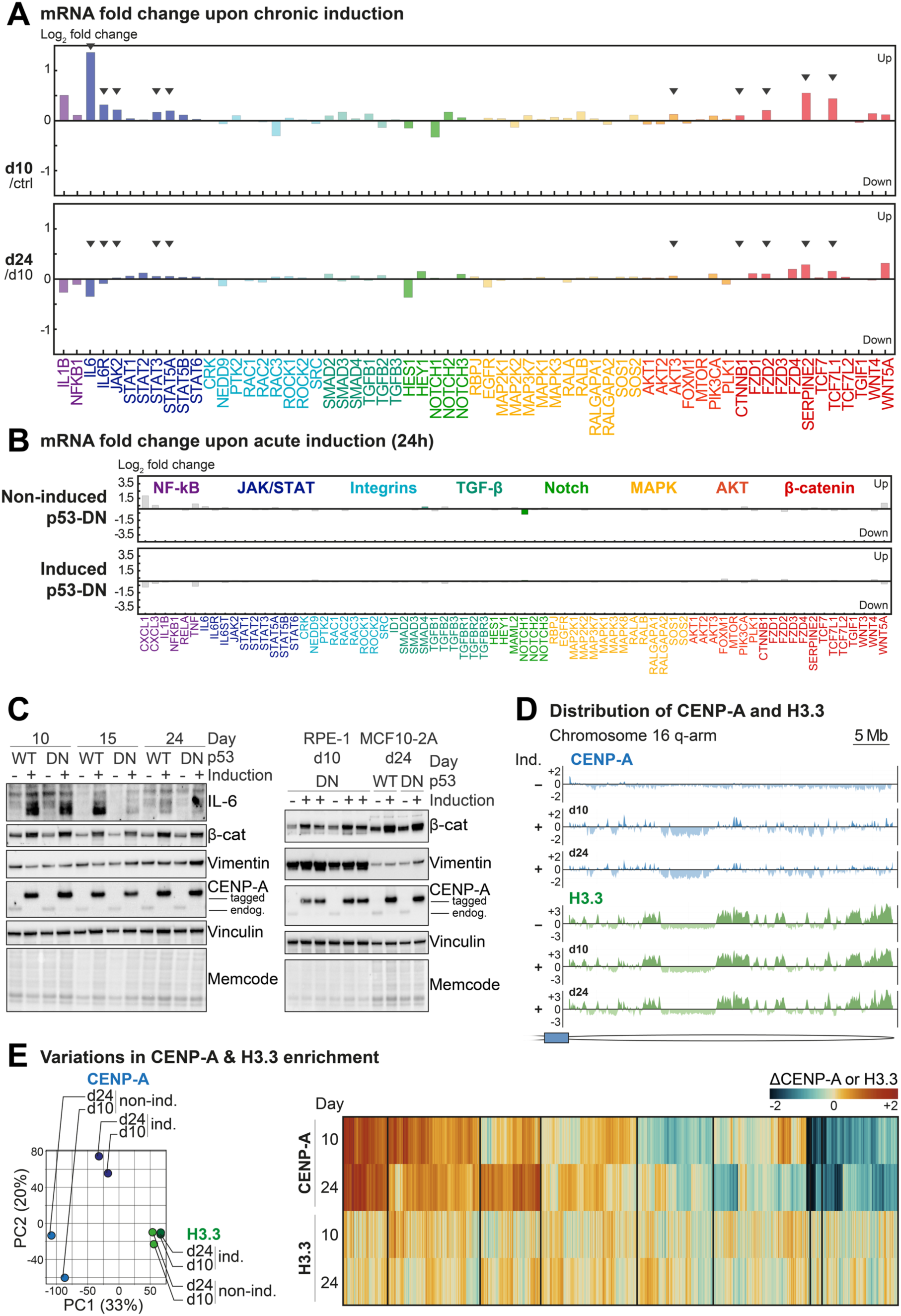
Impact of CENP-A on transcription and H3.3 and CENP-A localization. (A) mRNA expression of EMT genes in MCF10-2A cells 24 days post-induction in a biologically independent experiment (related to Figure 3). Bar plots show log_2_ fold change (TMM-normalized) in the pseudo-bulk RNA dataset used in Figure 4. Genes are grouped by signaling pathway and functional category (inflammation, left, or development, right). Arrows denote strong and consistent behavior with that observed in Figure 3A. (B) mRNA expression of EMT genes in MCF10-2A cells 24h post-induction. Bar plots show log_2_ fold change relative to the p53-WT non-induced condition (TMM-normalized). Genes are grouped by signaling pathway and functional category (inflammation, left, or development, right). Colored bars indicate statistically significant changes (adjusted p-value < 0.05). (C) Western blot analysis of total protein extracts from MCF10-2A or RPE-1 cells (-Dox or +Dox), with vinculin as loading control. Primary antibodies are indicated on the right. *IL-6 detection as multiple bands, consistent with its presence as differentially modified (glycosylated) isoforms, with apparent molecular weights from 23 to 30 kDa. **Non-phosphorylated β-Catenin (stable/active form) recognizing the functionally active protein (cell-cell adhesion and canonical Wnt signaling). (D) Genomic distribution of CENP-A (top, blue tracks) and H3.3 (bottom, green tracks) in cells with baseline or elevated CENP-A levels. Tracks show enrichment relative to input (counts per million, z-transformed log_2_ ratio) at bins of 100 kb. Enriched and depleted bins are highlighted in different colors (darker shades for enriched and lighter shades for depleted bins). (E) Variations in CENP-A and H3.3 enrichment. *Left:* Principal component analysis (PCA) of all ChIP-seq samples. *Right:* Complete heatmap of CENP-A and H3.3 enrichment at gene TSS (related to Figure 3D).

**Figure S4.**
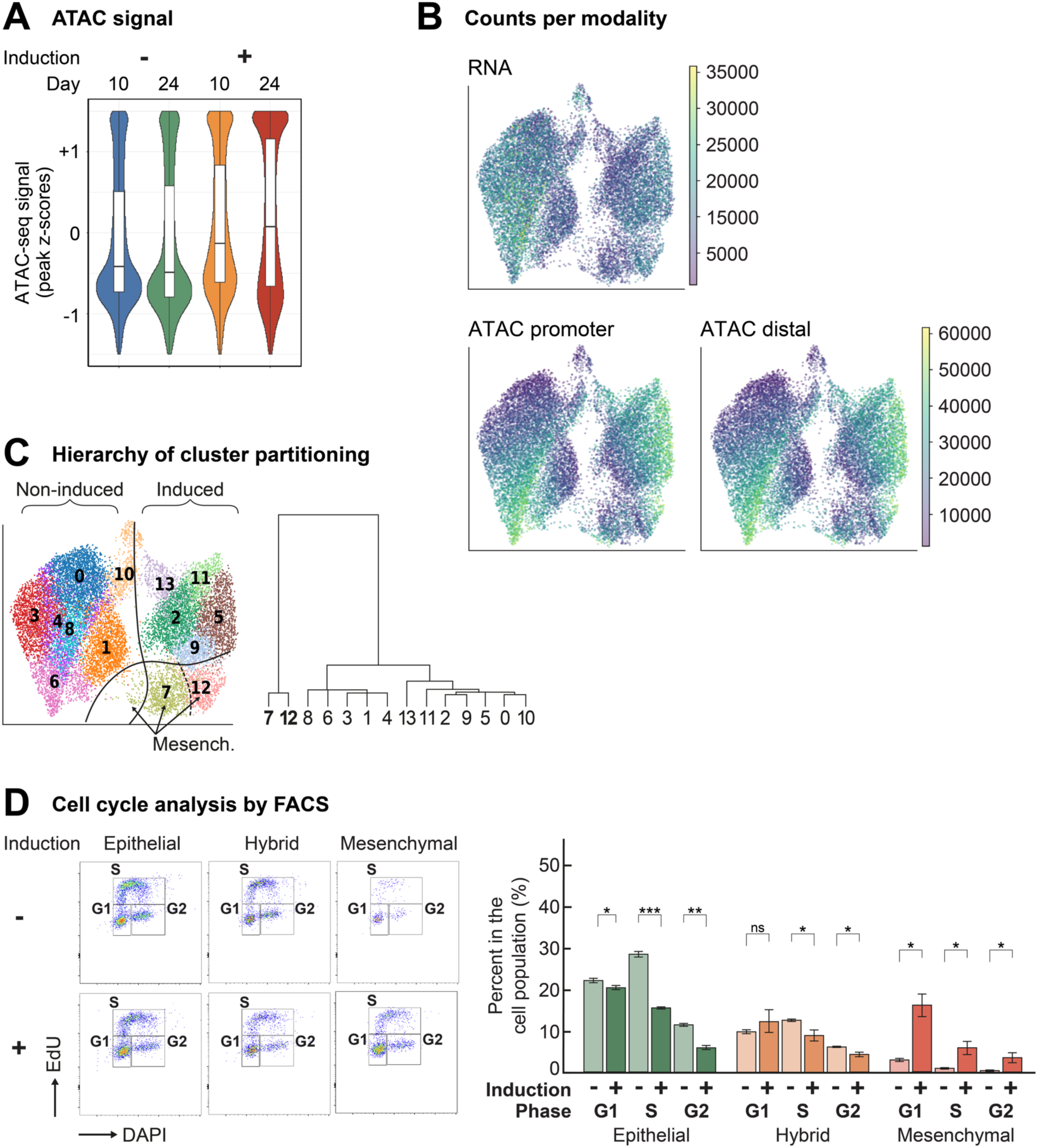
Impact of CENP-A on chromatin and cell cycle. (A) UMAPs of Mowgli embeddings showing counts per modality (RNA and ATAC). (B) Distribution of z-score normalized ATAC peak signals (same peaks across all samples). (C) Dendrogram showing hierarchical partitioning of Leiden clusters (related to Figure 4). (D) Cell cycle analysis. *Top:* FACS profiles at day 14 showing cells in “G1”, “S” and “G2” phases among epithelial, hybrid and mesenchymal cell populations, based on EdU and DAPI intensities. *Bottom:* Percentages based on FACS analysis. Plots show mean ± standard deviation for 2 biologically independent experiments. Statistical significance tested by two-tailed Welch’s t test. * = p-value < 0.05, ** = p-value < 0.01, *** = p-value < 0.001.

**Figure S5.**
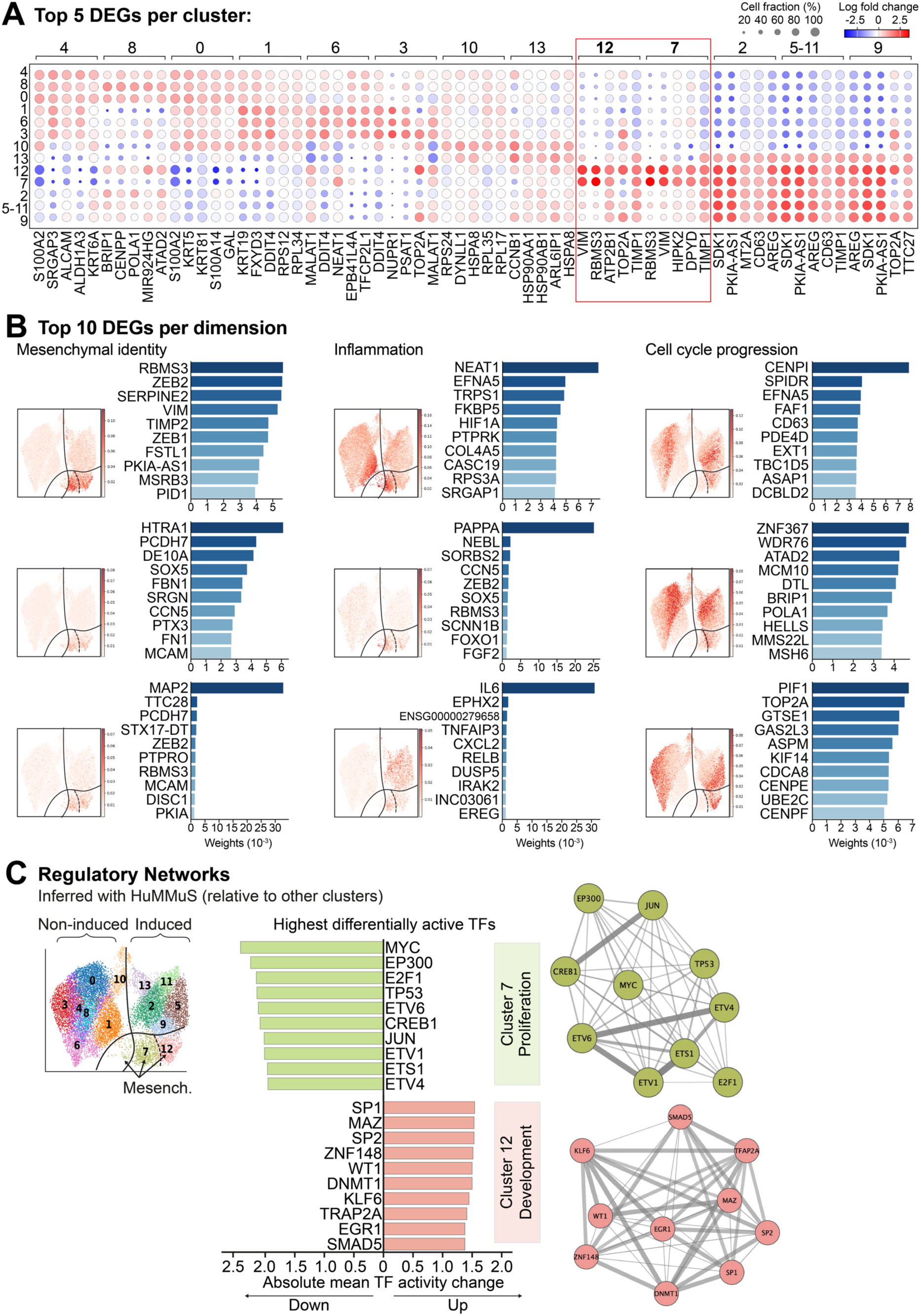
Gene regulatory analysis of mesenchymal clusters in the single-nucleus multiomics. (A) Top 5 DEGs genes per cluster. (B) Top 10 genes per Mowgli dimension. UMAPs show Mowgli’s embedding colored by factor weight. Bar plots display the top genes contributing to each factor, ranked by weight (RNA modality). Asterisks denote markers relevant to the corresponding biological aspect (mesenchymal identity, inflammation or cell cycle progression). (C) *Left:* Differential activity analysis (relative to all the other clusters) showing the top 10 highest differentially active Transcription Factors (TFs) per cluster. *Right:* TF cooperation networks. Lines width according to the strength of the connection (based on the number of shared targets across each TF pair).

**Figure S6.**
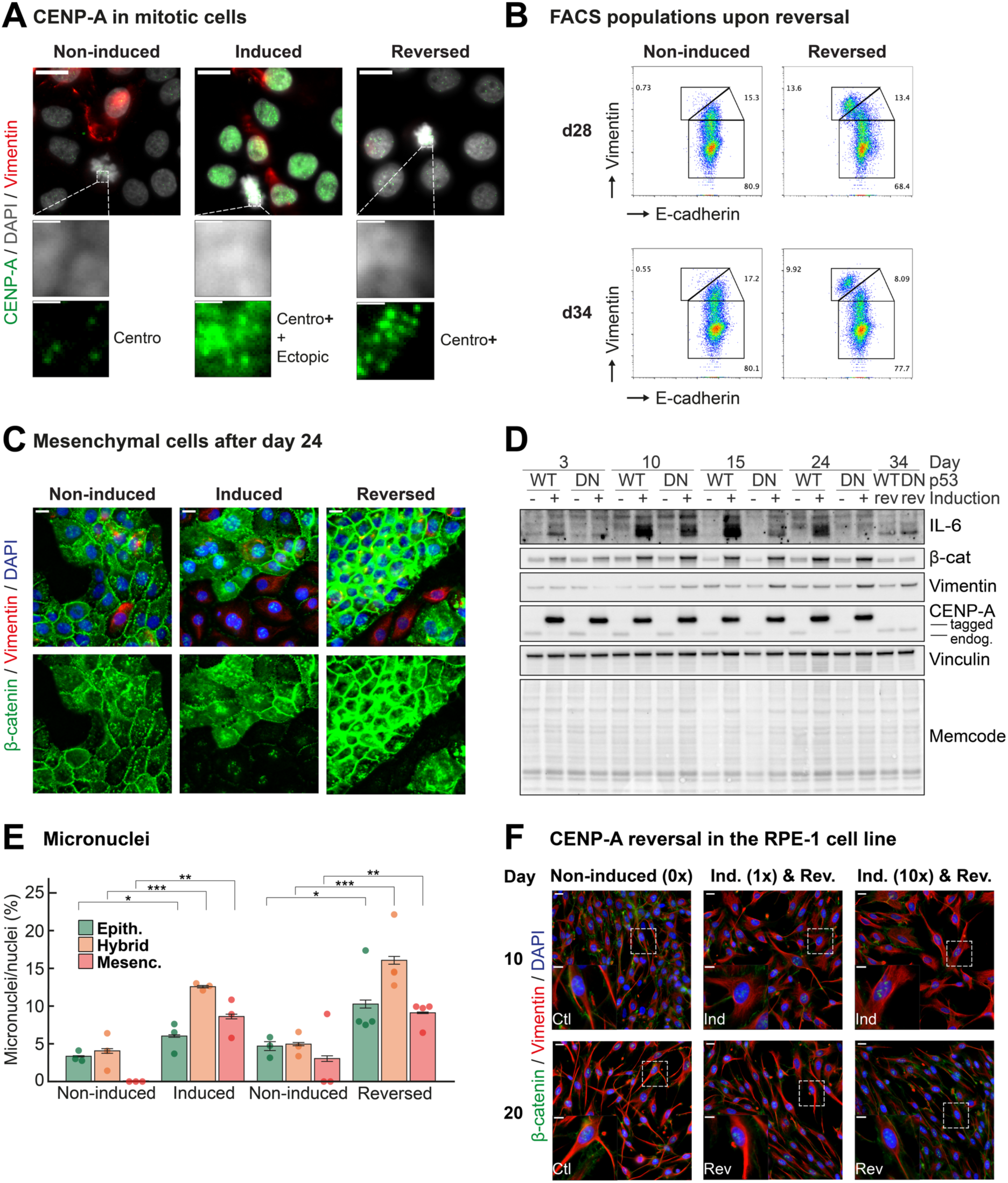
Cell populations and CENP-A localization upon reversal to basal CENP-A levels. (A) CENP-A localization in mitotic cells. *Top:* Representative epifluorescence images of cells immunostained for CENP-A (green) and Vimentin (red), with DNA counterstained with DAPI (grey). Scale bars = 20 µm. Squared boxes indicate cells on the bottom. *Bottom:* Insets of representative chromosome arms. Scale bars = 20 µm (wide fields), 2 µm (insets). (B) FACS profiles post-reversal (compared to induction in Figure S1D), at days 28 and 34, showing epithelial (E); hybrid (H) and mesenchymal (M) states based on vimentin and E-cadherin expression levels (relative to FACS analysis at day 24 from Figure S1). (C) Cell morphology and β-catenin localization in p53-DN MCF10-2A cells at day 34. Epifluorescence images showing β-catenin (green), vimentin (red) and DAPI (blue). Scale bars = 20 µm. (D) Western blot analysis of total protein extracts from MCF10-2A cells (-Dox or +Dox), with vinculin as loading control. Primary antibodies are indicated on the right. *IL-6 detection as multiple bands, consistent with its presence as differentially modified (glycosylated) isoforms, with apparent molecular weights from 23 to 30 kDa. **Non-phosphorylated β-Catenin (stable/active form) recognizing the functionally active protein (cell-cell adhesion and canonical Wnt signaling). (E) Quantification of micronuclei for each EMT state in control and CENP-A induced-cells (day 24) and upon reversal to basal CENP-A levels (days 27-34). Plots show mean ± 95% confidence interval. Dots indicate mean per experiment. n > 400 nuclei/state across ≥ 3 fields for each replicate. N = 3 or 4 biologically independent experiments. Statistical significance tested by two-tailed Welch’s t-test. * = p-value < 0.05, ** = p-value < 0.01, *** = p-value < 0.001. (F) Impact of CENP-A induction and reversal on the RPE-1 cell line. Epifluorescence images show β-catenin (green), vimentin (red), and DAPI (blue). Scale bars = 20 µm. Squared boxes indicate cells in the bottom-left corners (insets). Scale bars = 20 µm (wide fields), 10 µm (insets).

